# Pathogen cell wall degrading enzymes facilitate extracellular vesicles to deliver RNA into plants

**DOI:** 10.64898/2026.08.07.743426

**Authors:** Lorenz Oberkofler, Constance Tisserant, Chantal Krüger, Monica Rodriguez-Rendon, Charlotte Seydel, Andrea Cheradil, Nassim Safari, Ali Biabani, An-Po Cheng, Steffen Ostendorp, Andreas Klingl, Julia Kehr, Silke Robatzek, Arne Weiberg

## Abstract

Extracellular vesicles (EVs) can deliver RNA and proteins into host cells to manipulate immunity; but how EVs transverse the cell wall is unknown. Using the fungal pathogen *Botrytis cinerea* that induces cross-kingdom RNA interference in plants, we uncovered that EV-mediated RNA delivery was dependent on cell wall degrading enzymes. Through fluorescence and transmission electron microscopy, and molecular genetic techniques, we demonstrate that EV-associated proteins compromise the plant cell wall, thereby facilitating RNA delivery. Notably, cell wall degrading enzymes are commonly associated with EVs across plant-colonizing bacterial, fungal and oomycete species, indicating a conserved role in EV transport across cell walls. These findings uncover a formerly unknown mechanism by which cell wall degrading enzymes facilitate EVs to transverse the cell wall for cargo delivery in cross-kingdom communication.

## Introduction

Microbial pathogens such as bacteria, fungi and oomycetes deliver RNA and protein effectors into host cells to modulate plant innate immunity (1). In recent years, extracellular vesicles (EVs) have been realized as mediators of intercellular and cross-kingdom communication that has become evident across all domains of life. Notably, the plant pathogenic fungus *Botrytis cinerea* releases EVs containing small RNAs (Bc-sRNAs) that suppress host plant immunity through cross-kingdom RNA interference (cross-kingdom RNAi); a mechanism that is conserved in bacteria, fungi, and oomycetes interacting with plant or animal hosts in parasitic or mutualistic ways (2–7). Before *B. cinerea* EVs are endocytosed by plant cells (7), their traversal through the plant cell wall composed of cellulose, hemicellulose, and pectin poses a key question, given EV size constraints (8). Here, we provide evidence that fungal-secreted cell wall degrading enzymes (CWDEs) associated with EVs facilitate the degradation of cell wall components, allowing delivery of RNA cargo to access plant cells.

## Results

### Small RNA delivery via *Botrytis* EVs depends on extravesicular compounds

Following a prior investigation (7), we employed a differential ultracentrifugation protocol to collect BcEVs within a pellet fraction (hereafter referred to as crude BcEVs) either from the supernatant of fungal liquid culture (Fig.S1) or from the plant apoplast in *B. cinerea*-infected *Arabidopsis thaliana* (Fig.S2). These BcEVs were found to contain known cross-kingdom Bc-sRNA effectors (5, 8) as detected by stem-loop reverse transcription (RT)-PCR (Fig.S1, Fig.S2) and Illumina sequencing (Fig.S3, Tab.S1). The Bc-sRNAs within these crude BcEV samples exhibited resistance against enzymatic degradation. Even after Triton X-100 treatment, BcEV nanoparticles displayed only partial disruption but not full degradation in TEM images. Bc-sRNAs within crude BcEV samples remained detectable upon MNase and Proteinase K incubation (Fig.S4), implying RNA encapsulation inside BcEVs and possible protection by RNA-binding proteins (9).

Utilizing a transgenic *A. thaliana* line expressing a GFP-based, switch-on cross-kingdom RNAi reporter (10) (Fig.S5), we demonstrated that infiltrated crude BcEVs could elicit cross-kingdom RNAi in leaves (Fig.1A-B). Furthermore, plants treated with these crude BcEVs exhibited downregulation of known *A. thaliana* target genes of Bc-sRNAs (5) (Fig.S6). Notably, this gene suppression was absent in an *atago1* mutant line, corroborating previous evidence that plant AGO1 mediates cross-kingdom RNAi (5). The crude BcEVs also triggered induction of the plant immune marker genes *Pathogenesis-Related (PR)1* and *Plant Defensin (PDF)1.2*, as well as the two immune receptor-like kinases, *Flagellin22-Induced Receptor-like Kinase (FRK)1* and *Botrytis-Induced Kinase (BIK)1* (Fig.S6).

**Figure 1:**
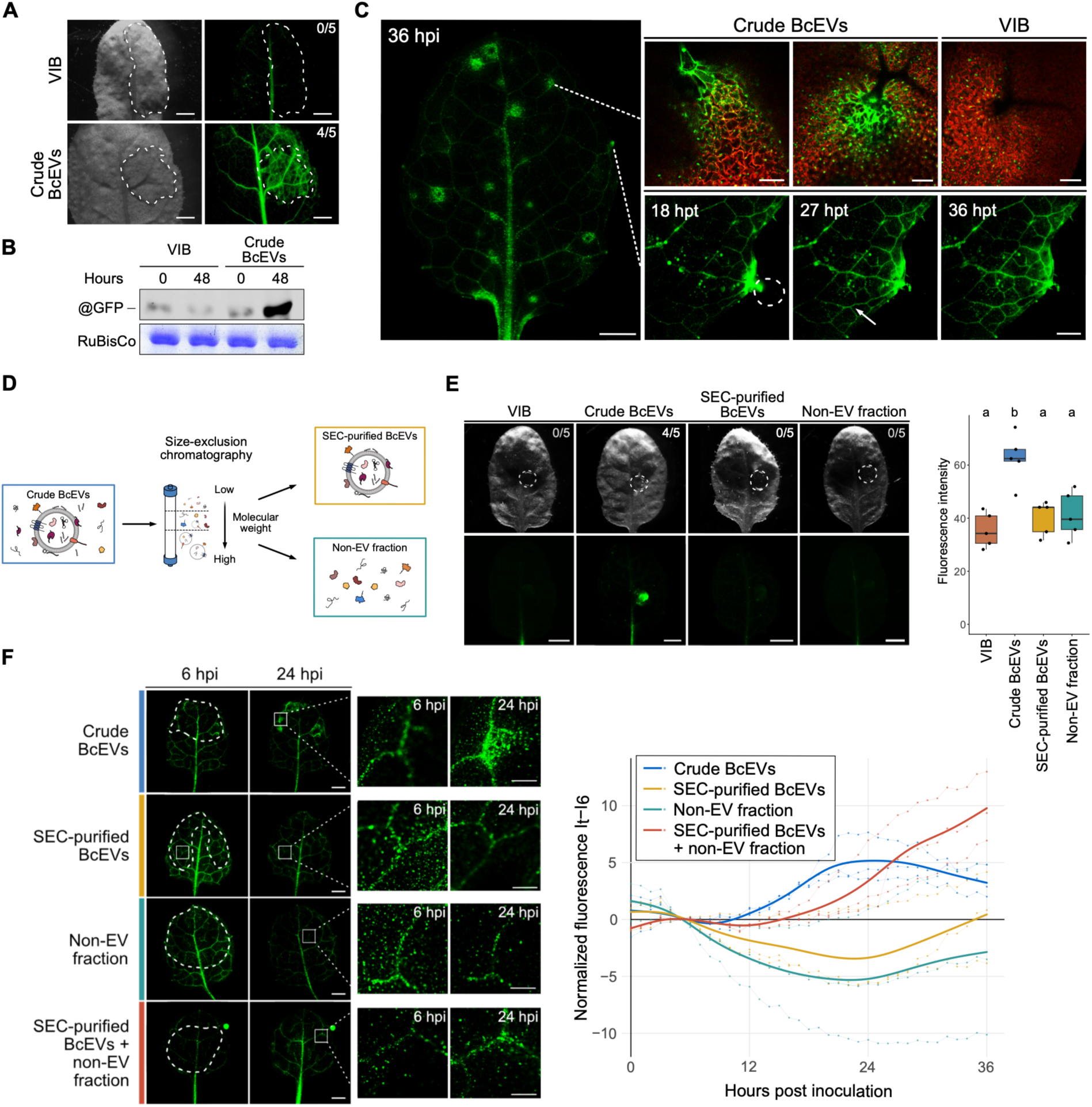
*Botrytis* EVs trigger cross-kingdom RNAi in plants. A) Crude BcEVs infiltrated into GFP reporter plant leaves at 24 hours post treatment, brightfield (BF, left) and GFP (right) images. Vesicle isolation buffer (VIB) was used as control. Dotted lines indicate area of infiltration. Scale bars represent 2 mm. B) Immunoblot analysis of GFP using total protein extracts of reporter plants upon BcEV or VIB treatment. Ribulose-1,5-bisphosphate carboxylase (RuBisCo) was used as loading control. C) Incubation with 50 µl crude BcEVs (1,1×10^13^ particles/ml) on the entire reporter plant leaf surface or 10 µl at the leaf edge in hydathode proximity (dotted circle). Arrow indicates GFP spreading in leaf veins. Scale bars indicate 2 mm for left image, 100 µm for top and 1 mm for bottom row. D) Schematic view of SEC-based BcEV purification. E) Local drop inoculation on GFP reporter plant leaves with 10 µl crude BcEVs or SEC-purified BcEVs (2,1×10^13^ particles/ml). Epifluorescence stereo-microscopy images show BF or GFP at 36 hpi. Numbers indicate events of GFP activation per total number of treated reporter leaves. Scale bars indicate 2 mm. Fluorescence intensity was quantified in inoculated areas. Statistical significance was determined by pairwise ANOVA followed by Tukey’s HSD; treatments not sharing the same letter are significantly different (p < 0.05). F) GFP reporter plant leaves were drop-inoculated with 5µl crude BcEVs (2,8×10^13^ particles/ml), SEC-purified BcEVs (9,1×10^13^ particles/ml), non-EV fraction, or SEC-purified BcEVs mixed with non-EV fractions of the same sample. Time-course fluorescence microscopy was carried out from 0 – 36 hours post inoculation and GFP signals were quantified on three independent leaves. Dashed white lines indicated area of BcEV treatment; the white square at 24 hpi marks a representative site of GFP activation, showed in higher magnification in adjacent panels. Scale bars in images represent 2 mm. GFP signals were imaged and quantified each 60 minutes. Normalized GFP fluorescence intensity (It – I6hrs) was plotted to quantify GFP activation. Dots connected by thin lines represent individual biological replicates (n = 3); bold lines represent mean values.

To exclude the possibility that plant stress responses activated the GFP reporter, we tested crude BcEVs derived from another *B. cinerea* strain, D08_H24, which does not produce the Bc-sRNAs recognized by the GFP reporter (8). Plants treated with D08_H24 crude BcEVs exhibited induction of the immunity marker *PDF1.2* (Fig.S7); however, even at a 5-fold higher concentration, these vesicles did not trigger GFP activation (Fig.S8, Fig.S9). This supports the conclusion that GFP reporter activation is specifically linked to Bc-sRNA delivery via BcEVs.

Applying crude BcEVs from *B. cinerea* strain B05.10 directly onto the leaf surface was sufficient to induce cross-kingdom RNAi (Fig.1C), suggesting possible natural entry pathways into the leaf interior. We propose two entry routes, likely involving the trichome basal cells and hydathodes.

GFP induction was observed to expand from the trichome base, indicating cell-to-cell transfer of Bc-sRNAs into neighboring epidermal cells (Fig.S10, Video S1). Entry through hydathodes was inferred from GFP activation and spread along vascular bundles (Fig.S10, Video S2).

Crude BcEV fractions might contain co-precipitates of soluble proteins and other low molecular weight molecules. To deplete such co-precipitates, we conducted BcEV purification though size exclusion chromatography (SEC) (11) (Fig.1D). Upon SEC purification, Bc-sRNAs persisted within the BcEV fraction but were undetectable in the non-EV fraction (Fig.S11). When GFP reporter plants were treated with SEC-purified BcEVs, cross-kingdom RNAi activity was abolished despite the continued presence of Bc-sRNAs (Fig.1E). Conversely, complementation experiments mixing SEC-purified BcEVs with the non-EV fraction from the same sample preparation demonstrated that Bc-sRNAs within purified BcEVs could still induce cross-kingdom RNAi (Fig.1F, Fig.S12). This combined treatment partially restored GFP activation, albeit with reduced intensity and a delayed response compared to crude BcEV treatment. These findings suggest that extravesicular compounds, either of protein or non-protein nature, present in crude BcEVs and the non-EV fraction are required for efficient Bc-sRNA delivery into host cells.

### Cell wall degrading enzymes associate with BcEVs

To determine whether the extravesicular compound(s) is of protein nature, we treated crude BcEVs either with Proteinase K or heat to inactivate solely the extravesicular proteins or all proteins in crude BcEV samples. Applying these treated crude BcEVs to GFP reporter plants confirmed the critical role of BcEV-associated proteins in mediating cross-kingdom RNAi (Fig.2A, Fig.S13). To ensure these treatments did not compromise BcEV integrity or Bc-sRNA cargo, we verified EV structure via TEM imaging (Fig.2B) and nanoparticle tracking analysis (NTA), and confirmed the presence of Bc-sRNAs through RT-PCR (Fig.S13).

**Figure 2:**
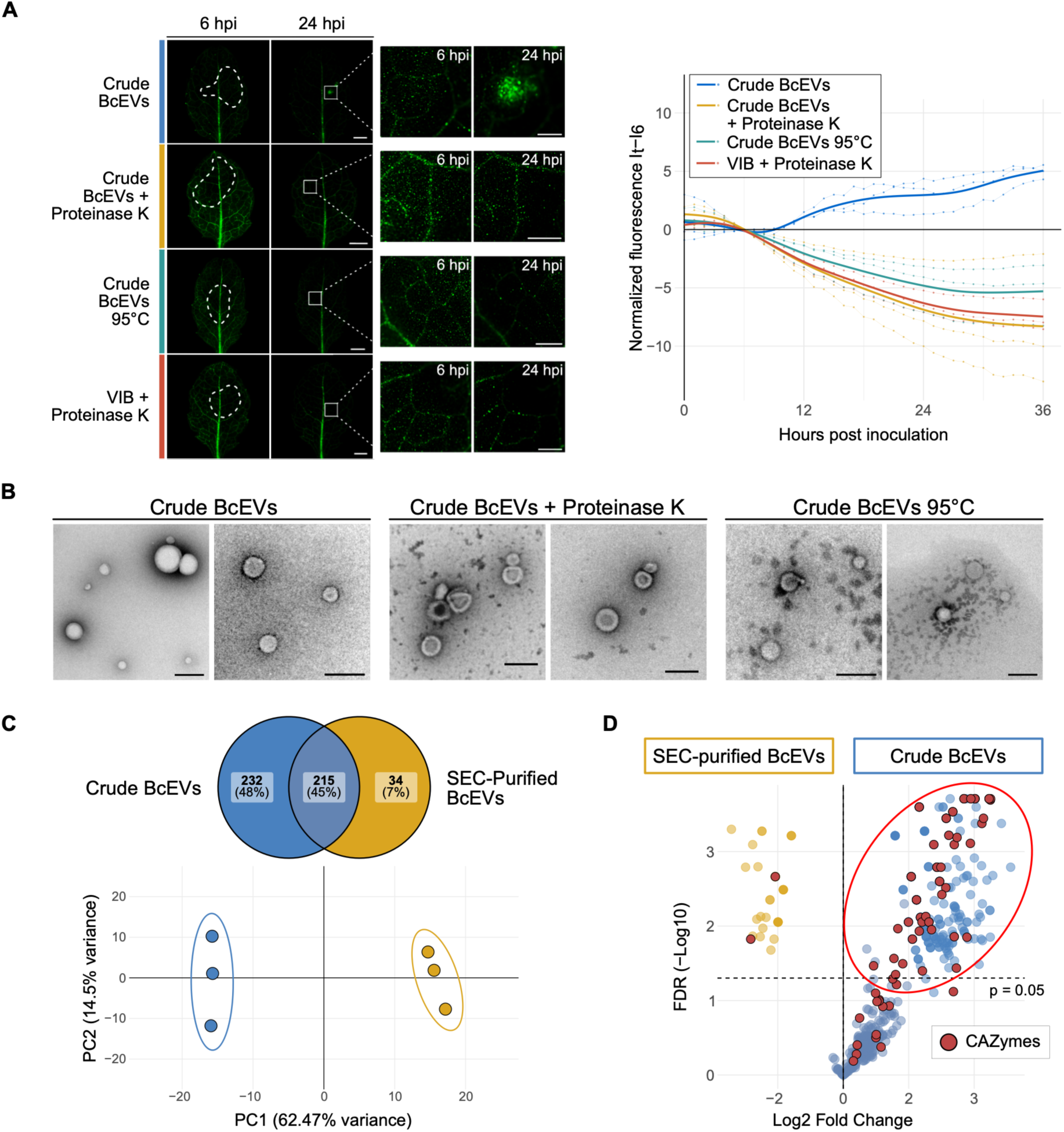
*Botrytis* EV-associated proteins are crucial for RNA delivery. A) GFP reporter plant leaves were drop-inoculated with 5 µl crude BcEVs (8,1×10^12^ particles/ml) without treatment, after treatment with proteinase K or after heat treatment at 95°C. Dashed white lines outline the area of BcEV treatment; the white square at 24 hpi marks a representative site of GFP activation, shown magnified in the adjacent panels. VIB buffer with proteinase K was used as control. Scale bars in images represent 2 mm. Normalized GFP fluorescence intensity (It – I6hrs) was plotted to quantify GFP activation. Dots connected by thin lines represent individual biological replicates (n = 3); bold lines represent mean values. B) TEM images of crude BcEVs before and after treatment with proteinase K or heated at 95°C. Scale bars indicate 200 nm. C) Principal component analysis of MS data identifying proteins in crude BcEV and SEC-purified BcEV samples. Data points represent three independent BcEV isolations. D) Volcano plot of protein hits identified in three biological replicates of crude BcEV and SEC-purified BcEV samples. The p-values were calculated based on a Benjamini-Hochberg corrected pairwise t-test.

To elucidate the protein composition of crude BcEVs, we next conducted a comparative proteomic analysis between crude BcEVs and SEC-purified BcEVs collected from fungal axenic culture supernatant using liquid chromatography-tandem mass spectrometry (LC-MS/MS). The results revealed largely distinct proteome profiles for the two BcEV samples, with over 50% of identified proteins being exclusively detected or significantly enriched in the crude BcEV fraction (Fig.2C, Tab.S2). Notably, overrepresented proteins in the crude BcEVs included putative signal peptides (SP) and predicted carbohydrate active enzymes (CAZymes). By contrast, SEC-purified BcEVs contained SP-free proteins and a higher proportion of predicted transmembrane proteins (Fig.S14, Fig.S15). Among the identified CAZymes, 69 potential cell wall degrading enzymes (CWDEs) were detected, with 45 significantly enriched in crude BcEVs, whereas only two were enriched in SEC-purified BcEVs (Fig.2D, Tab.S3). The CWDEs predominately belonged to glycoside hydrolases (GHs), with enzyme activities targeting diverse carbohydrate substrates typical of both fungal (chitin, glucan, mannan) and plant cell walls (cellulose, hemicellulose, pectin) (Fig.S16). Additionally, eleven putative adhesins - including proteins comprising carbohydrate-binding modules (CBM) and glycosylphosphatidylinositol (GPI)-anchor domains – were identified, suggesting potential roles in BcEV tethering to the plant surface (Tab.S2).

### *Botrytis* EV-associated proteins degrade the plant cell wall

Given the abundance of CWDEs in the crude BcEVs, we performed TEM imaging of leaf cross-sections at sites of GFP reporter activation following crude BcEV treatment. These sites exhibited a thinner, more ruptured plant cuticle compared to sections treated with SEC-purified BcEVs (Fig.3A). Furthermore, EVs were observed more frequently on the leaf surface after treatment with SEC-purified BcEVs, whereas fewer EVs were detected with crude BcEVs, suggesting a more efficient uptake of the latter into plant tissues.

**Figure 3:**
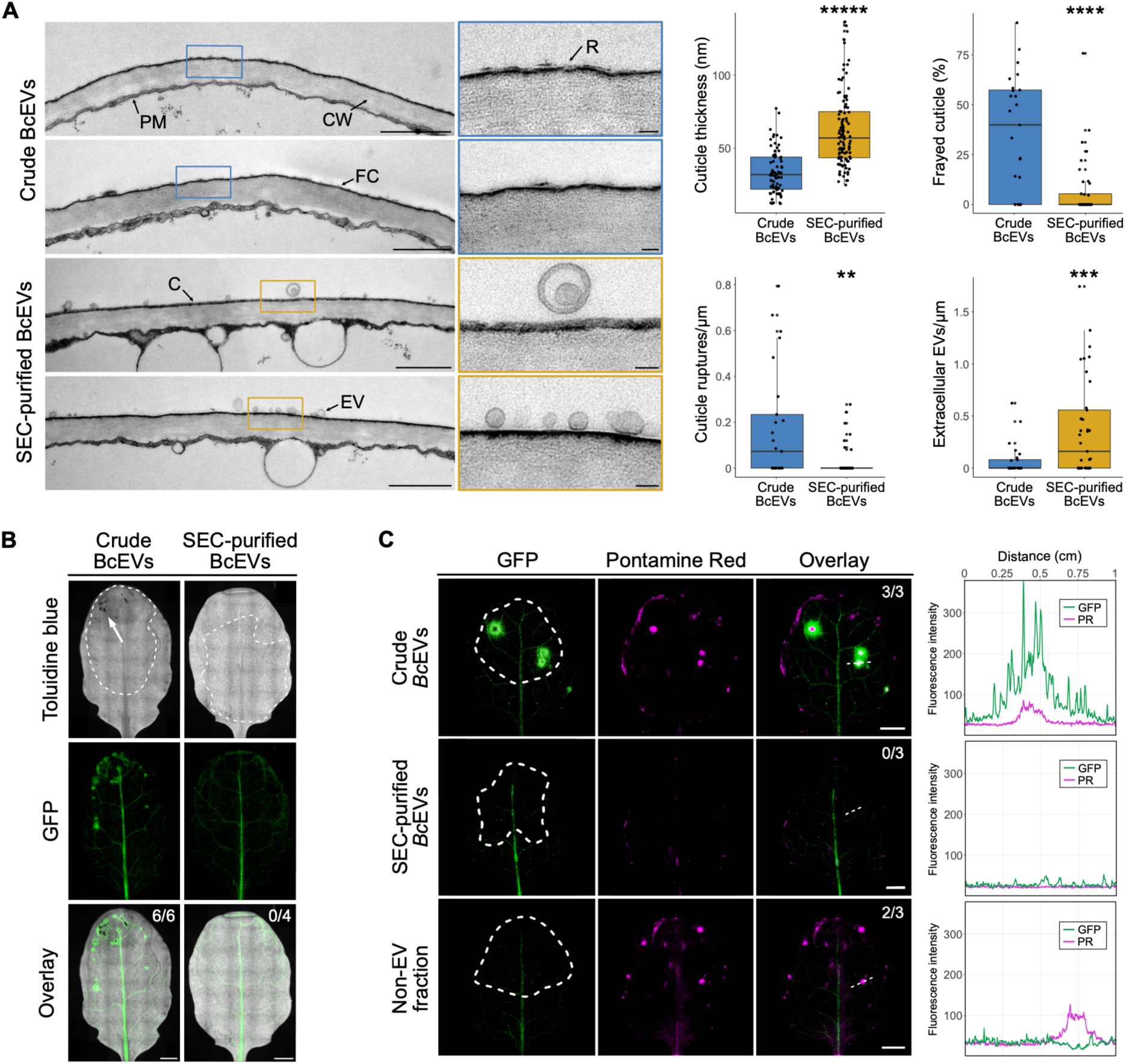
*Botrytis* EV-associated proteins damage the plant cell wall. A) TEM images of GFP reporter plant leaves treated either with crude BcEVs or SEC-purified BcEVs. Cross-sections were imaged at sites of GFP reporter activity. Plasma membrane (PM), cell wall (CW), cuticle (C), frayed cuticle (FC), cuticle rupture (R), extracellular vesicle (EV). Scale bars in images represent 1 µm in full and 100 nm in magnified images. Cuticle thickness and percentage of frayed cuticle was measured and cuticle ruptures as well as numbers BcEVs at the plant cell surface were counted across 61 images per treatment. Asterisks indicate significant differences using a pairwise t-test (p < 0.05). B) GFP reporter plant leaves were drop-inoculated either with crude BcEVs or SEC-purified BcEVs. Leaves were stained with toluidine blue and imaged in blue spectrum excitation and GFP channels. Numbers indicate GFP activation per total number of treated leaves. Dotted lines indicate areas of treatment. Scale bars represent 2 mm. C) GFP reporter plant leaves were drop-inoculated either crude BcEV, SEC-purified BcEV samples or the non-EV fraction. Leaves were stained with Pontamine fast scarlet 4B and imaged by fluorescence microscopy. Signal overlay was measured for GFP and Pontamine fast scarlet 4B fluorescence intensities. Dotted circles indicate areas of treatment and dotted straight lines indicate length of signal overlay analysis. Scale bars represent 1 mm.

To assess changes in cuticle permeability, toluidine blue staining was employed (12). The overlap of GFP reporter activity with toluidine blue signals in plants treated with crude BcEVs supported the hypothesis that these EVs compromised the cuticle, facilitating Bc-sRNA entry into plant cells (Fig.3B, Fig.S17). Further, staining with Pontamine fast scarlet 4B (Pontamine Red), which selectively labels accessible cell wall components like cellulose, revealed damage to the plant cell wall (13) – evident by overlapping fluorescence signals – upon crude BcEV treatment (Fig.3C, Fig.S17). Neither toluidine blue nor Pontamine Red signals were observed in plants treated with SEC-purified BcEVs. These findings collectively support the conclusion that CWDEs associated with crude BcEVs contribute to the disruption of the plant cuticle and cell wall, thus promoting efficient delivery of Bc-sRNAs into plant cells.

### Plant cell wall degradation is required for *Botrytis* EV-mediated RNA delivery

We next investigated whether the compromise of plant cell wall integrity is necessary to deliver Bc-sRNAs into plant cells via BcEVs. Treatment of reporter plants with SEC-purified BcEVs combined with Macerozyme (a commercial enzyme product primarily composed of pectinases and hemicellulases), which loosens the plant cell wall structure (Fig.S17), resulted in activation of the GFP reporter (Fig.4A, Fig.S18).

**Figure 4:**
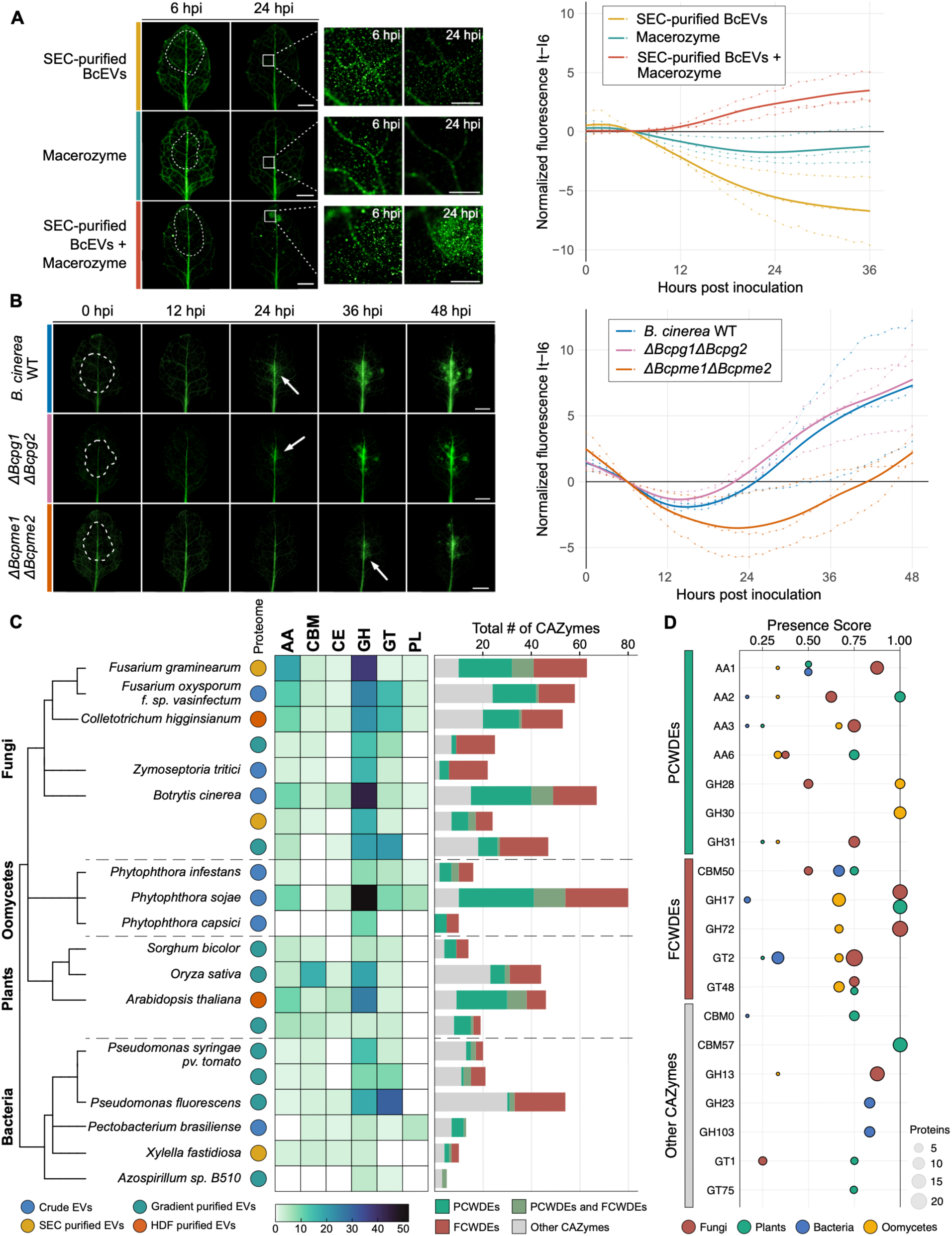
Plant cell wall degradation is required for *Botrytis* EV-mediated RNA delivery. A) Time point fluorescence imaging of GFP reporter plant leaves upon treatment with 5 µl SEC-purified BcEVs (5,7×10^13^ particles/ml) mixed with 0.5 % (w/v) Macerozyme. Macerozyme treatment without BcEVs was used as control. Dashed white lines outline the area of BcEV treatment; the white square at 24 hpi marks a representative site of GFP activation, shown magnified in the adjacent panels. Scale bars represent 2 mm. Normalized GFP fluorescence intensity (It – I6hrs) was plotted to quantify GFP activation. Dots connected by thin lines represent individual biological replicates (n = 3); bold lines represent mean values. B) Time point fluorescence imaging of GFP reporter plant leaves inoculated with *B. cinerea* WT, *ΔBcpg1ΔBcpg2* or *ΔBcpme1ΔBcpme2* ko mutants. Dotted lines in images indicate areas of treatment. Scale bars represent 2 mm. Normalized GFP fluorescence intensity (It – I6hrs) was plotted to quantify GFP activation. Dots connected by thin lines represent individual biological replicates (n = 3); bold lines represent mean values. C) Distribution and abundance analysis of CAZyme sub-families and CWDEs found in published EV protein databases of bacterial, fungal, oomycete and plant species. CAZyme sub-families are: auxiliary activities family enzymes (AA), carbohydrate-binding module family (CBM), carbohydrate esterase family (CE), glycosyl hydrolase (GH), glycosyl transferase family (GT), polysaccharide lyase family (PL). EV purification methods: blue circle refers to crude EV isolation by ultracentrifugation, yellow circle refers to size exclusion chromatography (SEC), green circle refers to low density-gradient fraction, red circle refers to high-density gradient fraction. D) Cross-kingdom abundance matrix of EV-associated CAZymes in bacterial, fungal, oomycete and plant species. Displayed families possess at least one kingdom and with a presence score >0.75.

Proteomic analyses highlighted an enrichment of pectin-degrading enzymes within the crude BcEV sample, including *B. cinerea* polygalacturonidase (BcPG)2 and pectin methylesterases (BcPME)1 and BcPME2. BcPMEs remove methyl groups from pectin - modifications commonly present in the plant cell wall – followed by BcPGs degrading the demethylated pectin (14). We hypothesized that BcPMEs and BcPGs facilitate Bc-sRNA delivery into plants via BcEVs. Genetic knockout (ko) mutants in these enzymes supported this model; the *ΔBcpg1ΔBcpg2* and *ΔBcpme1ΔBcpme2* mutant strains exhibited reduced virulence (Fig.S19), aligning with previous reports (15, 16). Both mutants possessed intact Bc-sRNA production (Fig.S19). Macerozyme rescued full infectivity of the *ΔBcpme1ΔBcpme2* strain, but not of *ΔBcpg1ΔBcpg2* (Fig.S20), suggesting that compromised Bc-EV-mediated RNA delivery is, at least in part, contributing to the reduced virulence of the PME mutant. In agreement, when inoculating GFP reporter plants, we observed that the *ΔBcpme1ΔBcpme2* strain caused delayed GFP activation compared to wild type (WT), whereas the *ΔBcpg1ΔBcpg2* mutant elicited GFP activation similar to WT (Fig.4B, Fig.S21). Comparable levels of fungal DNA were measured at 24 hours post inoculation - a time point when lower GFP reporter activation with *ΔBcpme1ΔBcpme2* was evident - suggesting that the reduced GFP response was not due to slower fungal colonization.

To further confirm a role of BcPMEs in EV-mediated Bc-sRNA delivery, we treated GFP reporter plants with crude BcEVs collected from WT and *ΔBcpme1ΔBcpme2* strains (Fig.S19). The mutant-derived vesicles occasionally activated the GFP reporter (two out of four replicates) whereas BcEVs from WT consistently triggered GFP signals. When treating with 1:3 diluted crude BcEVs, the *ΔBcpme1ΔBcpme2* mutant entirely failed to activate the GFP reporter, unlike WT (Fig.S22).

For co-localization of BcPMEs with BcEVs, we generated a *BcPME1-GFP* reporter strain using a Cas9-guided knock-in strategy. When inoculating plant tissue, the reporter strain indicated BcPME1-GFP accumulation in young mycelium at hyphal tips during plant infection (Fig.S23). Lipophilic staining confirmed co-localization of BcPME1-GFP with crude BcEVs but not with SEC-purified BcEVs (Fig.S24), consistent with the previous LC-MS/MS data. The presence of pectin-modifying enzymes in crude BcEVs was further supported by immunogold labeling using an anti-Pectinase antibody (Fig.S24). To assess the functionality of BcPME1 in BcEV-mediated RNA delivery, we tested if heterologous produced BcPME1 (Fig.S25) alone could restore Bc-sRNA delivery of SEC-purified BcEVs. Co-inoculating reporter plants with BcPME1 enzyme and SEC-purified BcEVs did not rescue GFP activation (Fig.S26).

Crude BcEVs also damaged plant cuticle integrity. Thus, we investigated whether cutinases may additionally contribute to BcEV-mediated Bc-sRNA delivery. The cutinase mutant strain *ΔBccutA* (17) and its isolated crude BcEVs were impaired in activating GFP in reporter plants (Fig.S27 and Fig.S28). Collectively, these results suggest that fungal PMEs and cutinases contribute to BcEV-mediated Bc-sRNA delivery into plant cells. Nonetheless, further fungal-secreted and BcEV-associated CWDEs likely act together to efficiently deliver Bc-sRNAs into plants for triggering cross-kingdom RNAi.

### CWDEs are a common feature of EV proteomes of plant-associated microbes

Plant pathogenic bacteria, fungi and oomycetes are well-known to secrete CWDEs to facilitate infection (18, 19). To explore whether these enzymes are frequently associated with EVs, we analyzed publicly available EV proteome datasets (Tab.S4). CWDEs were identified in six bacterial, nine fungal and three oomycete species, as well as two plant EV proteomes (Fig.4C, Tab.S5). It is important to note that these datasets were generated using various methods of EV isolation, fractionation, and purification. Consistent with our findings that CWDEs tend to be removed from BcEVs during purification, a similar reduction in CWDE levels were observed in purified EVs of the fungal pathogen *Colletotrichum higginsianum* and *A. thaliana* (20, 21).

Among all species examined, GHs were the most prevalent class of CWDEs similar to our observations in crude BcEVs. The presence of fungal CWDEs (FCWDEs) and plant CWDEs (PCWDEs) within EV proteomes suggests that these enzymes likely participate in cell wall remodeling during EV secretion. The GH17 family was consistently abundant across all eukaryotic species but was absent in bacteria. Oomycete EVs predominantly contained GH28 polygalacturonases and GH30 hemicellulases, whereas GH103 and GH23 enzymes were characteristic in bacterial EVs, both involved in peptidoglycan remodeling (Fig.4D, Fig.S29).

Additionally, numerous EV-associated proteins implicated in adhesion were identified. GPI-anchored members of the GH72 family were consistently abundant across all fungal EVs. Proteins with carbohydrate-binding modules (CBM) were also widespread, with LysM (CBM50) domains appearing across kingdoms, while malectin-like (CBM57) domains were restricted to plants. These binding motives are likely involved in tethering EVs to plant or fungal cell walls (22–24) and may also play roles in immune recognition through chitin sequestering and cell wall integrity sensing (25). Based on our EV proteome survey, we propose that carbohydrate-binding proteins and CWDEs are crucial for EV attachment and cargo delivery at cell-to-cell contact sites between plants and their associated microbes, facilitating communication and infection processes.

## Discussion

EVs are nano-sized carriers released by all living organisms for cell-to-cell and cross-kingdom communication, effectively transmitting functional cargo that influences organismal interactions. A fundamental question remains: how do EVs cross the cell wall barrier (26)? In this study, we demonstrate that CWDEs are associated with fungal EVs to facilitate the breakdown of complex plant cell walls, enabling efficient delivery of Bc-sRNAs. Consistent with this, regions of the plant where cell wall are naturally more permeable – such as trichome basal cells or hydathodes – may serve as accessible sites for EV exchange between plants and microbes (27, 28). Similarly, proteome analysis of EVs from various plant-associated microbes reveal that CWDEs are commonly present, suggesting a shared strategy to delivering cargo molecules into plant cells.

CWDEs associated with EVs could explain a targeted enzymatic attack at contact sites with the plant surface, enhancing cargo delivery efficiency. Furthermore, the composition of co-secreted proteins and enzymes associated with BcEVs is likely highly dynamic which makes vesicular transport capable of rapid adaptation to different target cells and organisms (29). This flexibility could allow BcEVs to adjust their cargo and enzymatic load for optimal interaction with plant or microbial species.

For successful cargo delivery, BcEVs must first adhere to the plant cell surface. In analogous systems, fungal pathogens deploy GPI-anchored glycoproteins (adhesins) to attach to plant cells for penetration (30). Supporting this, we identified CBM and GPI-anchored proteins within EV proteomes of *B. cinerea* and other plant-colonizing microbes. Though the precise mechanism involving CWDEs and CMBs in surface binding and cell wall remodeling requests further investigation.

While CWDEs are critical for Bc-sRNA delivery via BcEVs, their activity also produces cell wall breakdown products that can trigger plant immune responses known as damage-associated molecular pattern-triggered immunity (DAMP) (31). This aligns with our observation that BcEVs trigger immune-related gene induction in plants. We therefore propose a model in which BcEVs initially activate host immune responses while simultaneously delivering fungal sRNAs. By hijacking the plant RNA silencing machinery ultimately suppressing host immunity and facilitating pathogen infection.

Taken together, microbial EVs are increasingly recognized for their dual roles: inducing immunity and delivering effectors that suppress host defenses (32–34). In particular, *B. cinerea* and other pathogens utilize EVs to deliver sRNA and effector proteins into host plants modulating immunity in favor of infection (7, 35, 36). Conversely, plants deploy EVs carrying antimicrobial peptides and RNAs as part of their defense (6, 37, 38), highlighting EVs as central components in the ongoing host-pathogen arms race.

The delivery of bioactive molecules - such as RNA and peptides – via natural EVs or engineered lipid nanoparticles (LNPs) is now a prominent strategy in developing therapeutics, vaccines, and pesticides (39). Understanding how pathogen EVs utilize CWDEs and CMBs to facilitate specific and efficient cargo transfer could inform the design of advanced delivery systems, ultimately enabling targeted, biomolecule delivery with higher efficacy and specificity.

## Acknowledgments

We thank Matthias Hahn, Jan van Kan, Aline Voxeur, Luka Lelas for fungal strains. We are grateful to Adriana Hörmann, Su Cacace, Jennifer Grünert and Jessica Folgmann for technical assistance, Henning Kunz for access to plant growth facilities, as well as Andreas Brachmann for sequencing service. We also would like to thank Claude Becker and Arp Schnittger for proofreading and Martin Parniske, Caroline Gutjahr, Roger Innes, Niklas Schandry, Hannah McMillan, Alessa Ruf, Ben Koch, Santiago Priego-Cubero, Liza Rouyer, Eliana Mor, Elif Olkun, Bernhard Lederer and other members of the RU5116 and exRNA-PATH members for valuable discussions and suggestions.

## Author contributions

Conceptualization: LO and AW

Investigation: LO, CT, CK, MRR, CS, AC, AK, APC, SO, AB, NS

Writing – original draft: LO, CT, AW

Writing – review and editing: JK, SO, APC, CK, SR

Visualization: LO

Funding acquisition, project administration, supervision: AW

## Data and materials availability

Raw sequencing data of BcEV small RNAs are available at the NCBI-SRA archive under the BioProject: PRJNA1381111.

## Materials and Methods

### Fungal and plant materials

*Botrytis cinerea* wild type isolates B05.10 and D08_H24 (*40*) as well as *ΔBcpg1ΔBcpg2* (*41*), *ΔBcpme1ΔBcpme2* (*42*), *ΔBclip1* (*43*) and *ΔBccutA* (*44*) mutants were used in this study. For conidiospore production, all fungal strains were grown on HA medium (10 g/l mal extract,4 g/l yeast extract, 4 g/l glucose, 15 g/l agar) at room temperature under constant near ultraviolet light exposure for 10 to 14 days. For BcEVs isolation, 200 ml water inoculated with 5×10^6^ conidia per ml was added to 800 ml liquid minimal medium (LMM: 20 g/l sucrose, 4 g/l NaNO3, 1 g/l K2HPO4, 0.5 g/l KCl, 0.5 g/l MgSO4 and 0.01 g/l FeSO4; pH adjusted to 6.0) and cultures were incubated under low shaking condition (100 rpm) at room temperature for 3 days. *Arabidopsis thaliana* WT, cross-kingdom RNAi reporter plants (*45*) and *atago1-27* mutants (*46*), all in the ecotype Col-0 background, were grown in soil under short day conditions (10 hours light/14 hours dark, 22 °C and 60 % relative humidity).

### Plant inoculation with *B. cinerea*

Infection assays with *A. thaliana* were performed on detached leaves from 4- to 5-week-old plants. *B. cinerea* conidia were resuspended in 1% malt extract at a final concentration of 2×10^5^ conidia/ml. 15 μl conidia suspension was dropped at the center of each leaf and inoculated in a humidity plastic box. Infected leaves were photographed at 72 hours post inoculation (hpi) and lesion area was measured using the Fiji software (ImageJ version 2.16.0). 0.5% (w/v) Macerozyme (Macerozyme R-10, Yakult Pharmaceuticals) was added to the conidiospore suspension in mutant infection rescue experiments.

For time course experiments with GFP reporter plants, detached leaves from 3-week-old *A. thaliana* seedlings were inoculated with 2×10^5^ conidia/ml resuspended in 1% malt extract medium pre-incubated for 1 hour. The pre-incubated conidiospore suspension was washed with sterile water before 5 μl was dropped at the center of the leaves and covered with a cover slide.

Growth rate of *B. cinerea* WT and mutant strains was recorded by placing an agar plug containing young mycelium in the center of a HA agar containing Petri dish. To test the impact of Macerozyme of fungal growth, Macerozyme was added with a final concentration of 0.5% (w/v) to a spore suspension (2×10^4^ conidia/ml) and 10 µl of the suspension were pipetted on a HA agar containing Petri dish. Plates for all experiments were incubated for 5 days at room temperature under constant near ultraviolet light exposure. Mycelial growth was determined by measuring the radial growth of colonies every 24 hours.

### Extracellular vesicle isolation and analysis

To collect crude BcEVs, fungal liquid cultures were centrifuged at 4,000 g for 20 min at 4°C. The supernatant was filtered through two layers of Miracloth filter paper (pore size 22-25 μm, Merck Millipore). Cellular debris was removed by centrifugation at 13.000 g for 50 min at 4 °C. Low-speed centrifugation steps were carried out using a fixed rotor (JA-10 rotor, Avanti J-26S XP centrifuge, Beckman Coulter).

The supernatant was collected and filtered through 0.22 μm sterile filters (Steritop, Merck). Crude BcEVs were collected by ultracentrifugation with a swing rotor (SW 32 Ti Rotor, Optima XE-90 Centrifuge, Beckman Coulter) at 100.000 g for 90 min, 4°C. Pellets from all tubes were pooled into one, the tube was filled with VIB buffer (20 mM Mes Hydrate, 2 mM CaCl2, 0.1 M NaCl, pH 6.0) (*47*) and EVs were reconcentrated by an additional centrifugation step, as described above. Crude BcEVs were resuspended in sterile-filtered VIB buffer and transferred to a protein LoBind reaction tube (Eppendorf, 88379). The crude BcEVs were kept on ice and used immediately for experiments. BcEVs were snap frozen in liquid nitrogen and stored up to a week at −80° C only for stem-loop RT-PCR experiments. The EV isolation pipeline was carried out in a cold room at 4°C and/or on ice.

To collect EVs from the AWF of *B. cinerea*-infected *A. thaliana*, 100 infected and 25 non-infected six-week-old plants were grown per experiment. Not infected *A. thaliana* was sprayed with 12 ml of 1% malt extract and *B. cinerea*-infected plants were sprayed with a conidia suspension (3.5 × 10^6^ conidia/ml) in 1% malt extract. After 4 days, the AWF was collected following the protocol of Rutter *et al*. (*47*).

For SEC purification, crude BcEV pellets were resuspended in 2 mL filtered VIB buffer and further purified using qEV2 iZON SEC-columns (iZon qEV2 columns, 70 nm series, IC2-70). Prior to EV loading, columns were flushed and equilibrated with filtered VIB buffer following manufacturer’s instructions. Fractions containing purified BcEVs (2-12 ml) and non-EV associated proteins (14-50 ml) were collected, enriched by an additional ultracentrifugation step (100.000 g for 90 min, 4°C) and resuspended in 100 µl filtered VIB.

### Lipophilic staining of EVs

For lipophilic staining, 50 µL of BcEVs were incubated with 1 mM FM4-64 (Invitrogen, T13320) at 4 °C for 1 hour, and imaged on a Leica Thunder imager using a 100×/1.44 OIL UV objective.

### Nanoparticle tracking analysis of EVs

For BcEV characterization using a nanoparticle tracking analyzer (NTA, ZetaView, Particle Metrix), vesicle samples were diluted to a concentration which resulted in approx. 200–400 particles per window. Measurements were performed at a cell temperature of 22°C, with vesicle size profiles and concentrations determined as the average of 11 camera positions across three independent replicates per sample.

### Transmission electron microscopy of EVs

Transmission electron microscopy (TEM) of BcEVs, 10 µl of BcEV samples were applied to 400 mesh carbon-coated copper grids and negative stained with 1% uranyl acetate as described previously (*48*). TEM was carried out with a Zeiss EM912 TEM (Carl Zeiss AG, Oberkochen, Germany), equipped with a 2k x 2k slow-scan CCD Camera (TRS, Tröndle Restlichtverstärkersysteme, Moorenweis, Germany) and operated at 80 kV and 120 kV in zero-loss mode, or a JEOL F200 cryo-(S)TEM (JEOL, Freising, Germany) with a XAROSA 20 megapixel CMOS camera (EMSIS, Münster, Germany) and operated at 200 kV.

### BcEV treatment and complementation assays

BcEVs samples were divided equally into three to four aliquots. Triton X-100 (Sigma-Aldrich, T8787-250ML) treatment was carried out for 5 min at 70°C with a final concentration of 0,5% or 1%. Proteinase K (Thermo Scientific, EO0491) was added to reach 50 μg/ml and incubated for 15 min at 37°C, and inhibited by the addition of phenylmethylsulfonyl fluoride (PMSF) to a concentration of 100 mM and an incubation of 10 min at room temperature. RNA digestion was performed with 1000 units of MNase (NEB, M0247S) for 30 min at 37°C. TEM images were taken and corresponding stem-loop RT-PCR were performed after 30 min incubation on ice with 1% Triton X-100, according to He *et al.* (*49*).

For protein degradation assays, crude BcEVs were resuspended in 200 µl VIB and split in 3 samples. The first was kept on ice, the second was subjected to 100 µg/ml Proteinase K treatment (NEB, P81075) for 30 min at 37°C and the third was heat treated at 95°C for 5 min. For cell wall loosening assays, Maceroenzyme mix was added to VIB and 100 µl purified BcEVs to a final concentration of 0.5% (w/v).

Fraction complementation assays were carried out by combining SEC-purified BcEVs with the non-EV fraction in a 1:1 ratio. Crude BcEVs, SEC-purified BcEVs and non-EV fractions were equally mixed in a 1:1 ratio with filtered VIB. BcEV fractions were measured via NTA to normalize particle concentration.

### BcEV treatment of plants

BcEV infiltration experiments were performed on detached leaves from 4-week-old *A. thaliana* reporter plants. VIB and BcEVs were infiltrated with a plastic syringe from the abaxial site and infiltration area was gently marked with a marker on the adaxial site. Exclusively infiltrated plant material was sampled for RNA extraction.

BcEVs drop inoculation experiments were performed with detached leaves from 2- to 3-week-old

*A. thaliana* reporter seedlings. Leaves were drop inoculated with 5 µl of BcEVs and covered with a cover slide. Crude BcEV pellets were diluted in 200 μl VIB buffer for plant reporter assays, or noted otherwise. For vasculature treatments, BcEVs were dropped next to the leaf in proximity of a hydathode. BcEV quantities were measured and normalized via NTA to result in the same concentration in all treatments. Seedlings in both conidia and BcEV treatments were incubated in a square petri dish sealed with parafilm to maintain humidity.

### GFP switch-on cross-kingdom RNAi reporter construct

A GFP switch-on cross-kingdom RNAi reporter was recently established in a transgenic *A. thaliana* plant line (*45*). In brief, the GFP reporter exclusively responds to the translocation of Bc-sRNA3.1 and Bc-sRNA3.2 from *B*. *cinerea* into the plant host. The CRISPR-type RNA endonuclease Csy4 (*50*) is co-expressed with a GFP version that is fused to the Csy4 recognition motif at its N-terminus. Csy4 constantly suppresses expression of the *GFP*, unless GFP expression is activated when Bc-sRNAs silence *Csy4*. A schematic view of the reporter construct is given in Fig.S5. The GFP reporter plant line allows to visualize *B*. *cinerea*-induced cross-kingdom RNAi in infected leaf tissue by fluorescent microscopy.

### Transmission electron microscopy of plant leaves

For TEM analysis of seedling epidermis after BcEV treatment, areas of leaves with GFP reporter signal were selected for sampling. Leaf material from the sampling area was cut in 1 mm^2^ pieces in fixation buffer (75 mM cacodylate, 2 mM MgCl2, pH 7.0) supplemented with 2.5% glutaraldehyde. Fixation was carried out as described previously (*51*). After several washing steps, post-fixation in 1% OsO4 in fixation buffer and en-bloc contrasting with 1% uranyl acetate was conducted. Samples were dehydrated in a graded acetone series and subsequently embedded in Spurr’s resin of medium rigidity (*52*). Ultrathin sections of approximately 60 nm were prepared and contrasted with lead citate (*53*). TEM was conducted with the Zeiss EM 912 equipped with an Omega-filter, operated at 80 kV in zero-loss mode (Carl Zeiss AG, Oberkochen, Germany) and a 2k x 2k slow-scan CCD camera (TRS Tröndle Restlichtverstärkersysteme, Moorenweis, Germany). Quantification of thickness, frayed and ruptured cuticle phenotypes as well as BcEV amount outside of the cuticle was done across 4 blocks for both conditions, with a total of 25 sections analyzed for crude and 36 for purified BcEV-treated leaves. Measurements were carried out on Fiji (version: 2.16.0) and results were normalized based on cuticle length determined on each section.

### Immunogold labelling of pectinases

For immunogold staining, 10 µl of BcEV sample was applied to 200 mesh nickel grids that were coated with collodium and a layer of approximately 10 nm carbon. After 5 min incubation, the sample was blotted from the grid with filter paper. Blocking was carried out twice for 5 min and 20 min with 0.1% bovine serum albumin (BSA) in 1x PBS buffer. The primary antibody (Pectinase polyclonal antibody, Bioss, BS-4550R) was applied to the grids for 30 min at a dilution of 1:400, then washed three times with blocking solution. The secondary antibody (Anti-Rabbit IgG whole molecule gold antibody, Sigma-Aldrich, G7277), was applied for 45 min at a dilution of 1:100, followed by two consecutive washing steps with 0.1% BSA, two washing steps with 1x PBS, and two washing steps with water. Directly after immunolabelling, the grids were negative stained for 2 min with 1% uranyl acetate and dried after blotting. Gold particles were counted in 10 images for both conditions, including respective controls containing only the secondary antibody, and the average particle number per µm^2^ was calculated.

### Plant cell wall staining and cuticle permeability assay

BcEV-, non-EV fraction- or Macerozyme-treated (0.5% w/v) reporter plant leaves were sampled at 24 hours post treatment and were stained in the dark with 0.1% (w/v) Pontamine fast scarlet 4B (Sigma-Aldrich, 212490) solution for 30 min at room temperature and washed 3 times with ddH2O before imaging. Cuticle permeability was detected by leaf immersion in 0.05% (w/v) toluidine blue (Roth, 0300.2) solution for 5 min, followed by three washes in ddH2O.

### Quantitative fluorescence microscopy

For time course experiments on a Thunder Imager (Leica), squared petri dishes containing treated reporter seedling leaves, were mounted on the Leica DMi8 Thunder Imager equipped with a QUANTUM high speed stage, a Leica DFC9000 GT camera and LED5 laser. Whole leaf images were captured with a 5x objective every 20 to 60 min (depending on experiment) for up to 48 hours. Fluorescent signals of GFP (Ex: 475 nm; Em: 506-535 nm), FM4-64 (Ex: 575 nm; Em: 641-642 nm), Direct Red 20 (Ex: 555 nm; Em: 578-590 nm) and Toulidine Blue (Ex: 390 nm; Em: 420-460 nm) were measured. Raw imaging data were processed using the Leica LAS X software (version: 3.9.1.28433) and GFP fluorescence intensity was quantified at inoculation sites, as indicated by the white dotted line, for all experiments involving conidiospore and undiluted BcEV treatments. For diluted BcEV treatments, fluorescence measurements were restricted to the enlarged region of interest, as the response was too subtle to be reliably quantified across the entire inoculation area. Measurements were carried out in Fiji (version: 2.16.0). For data analysis, mean grey values were normalized at given time points (It) by subtracting GFP intensity at 6 hpi (I6). Video editing was performed using Adobe Premiere Pro CC (version 13.0). Overview pictures of reporter leaves were taken using a M165 FC epifluorescence stereomicroscope (Leica) in bright field and with a GFP filter.

### Confocal microscopy

High resolution confocal microscopy was performed using the Stellaris 5 Confocal Laser Scanning Microscope (Leica), equipped with a 405 nm diode and a supercontinuum White Light Laser (WLL), with fluorescence detection using a Power Hybrid Detector HyDS. Images of EV treated reporter leaves were taken with a 20x objective. Fluorescent signals of GFP (Ex: 475 nm; Em: 509-535 nm) and chlorophyll A (Ex: 440 nm; Em: 670-685 nm) were captured.

### GFP immunoblot analysis

Imunoblot analysis of GFP expressed in *A. thaliana* was performed, as previously described (*45*). Proteins were separated on an 8% SDS-polyacrylamide gel at 80 volts for 30 min and 140 volts for 2 hours and transferred to PVDF membrane (Immobilon-FL) overnight at 4°C. Transferred membranes were blocked with 10 ml of 5% (v/v) skim fat milk in 1× PBS at 4°C for 1h on a rolling shaker. Membranes were incubated overnight with primary GFP Rabbit Polyclonal Antibody (600-401-215, Rockland). The membranes were incubated with secondary antibody α-rat IRdye800 (LI-COR) for 1 hour. Protein signals were detected using Odyssey imaging system (LI-COR).

For protein detection of BcPME1-GFP, *B. cinerea* cultures were grown in pectin-containing minimal medium consisting of Gamborg B5 supplemented with 10 mM KH₂PO₄, 0.5% pectin, and 2.5 mM glucose. Mycelial samples were collected after 28 hours, and culture supernatants were collected to analyze the secretome. Mycelial proteins were extracted in buffer containing 50 mM Tris-HCl, 150 mM NaCl, 1% Triton X-100 or NP-40, 10% glycerol, protease inhibitor cocktail, and PMSF. Supernatant proteins were concentrated using centrifugal filter units and subsequently precipitated with an equal volume of 20% TCA overnight at −20°C, followed by acetone washing. Protein samples were separated by SDS-PAGE, transferred to nitrocellulose membranes, and probed with a mouse anti-GFP antibody (JL-8, Takara/Clontech, 632381) followed by an HRP-conjugated anti-mouse secondary antibody (Sigma-Aldrich, A8924). Signals were detected by chemiluminescence using a ChemiDoc Touch imaging system (Bio-Rad). *B. cinerea* WT isolate B05.10 and free GFP samples were included as negative and positive controls, respectively.

### Stem-loop RT-PCR and qRT-PCR

Total RNA was extracted from *B. cinerea* mycelium, isolated BcEVs and *A. thaliana* plants using a CTAB-based method (*54*). Small RNA detection by stem-loop RT-PCR was carried out as described (*55*). Small RNA detection was performed using 1 µg of total RNA from mycelium and 100 µl of isolated BcEVs. Reverse transcriptase PCR products were separated on a 10% non-denaturing polyacrylamide gel followed by ethidium bromide staining. PCR products were purified from the gel and blunted by a Klenow reaction (NEB, M0212L) prior to cloning for Sanger sequencing.

For gene expression analysis, genomic DNA was removed by DNase I (ThermoFisher, EN0525) treatment following the manufacturer’s instruction. 1 μg of total RNA from each sample was used for cDNA synthesis using the Maxima H Minus reverse transcriptase (ThermoFisher, EP0752) and an oligo dT primer. Genomic DNA quantities and gene expression were measured by quantitative real-time PCR using the Primaquant low ROX qPCR master mix (Steinbrenner Laborsysteme GmBH). Differential gene expression level was calculated using the 2-ΔΔCt method (*56*). Primers used in quantitative and stem-loop RT-PCR are listed in Tab.S6.

### Small RNA sequencing

Total RNA extracts isolated from *B. cinerea* mycelium or BcEV samples were separated on a 15% polyacrylamide gel to purify sRNAs. Purified small RNAs were cloned for Illumina sequencing using the Next® Small RNA Prep kit (NEB) and sequenced on an Illumina HiSeq1500 platform. The sequencing raw data were pre-processed using the GALAXY Biostar server (*57*). Processed raw reads are deposited at the NCBI SRA server (BioProject PRJNA1381111). Raw reads were mapped to *B. cinerea* B05.10 reference genome (ASM14353v4) using the BOWTIE algorithm (Galaxy Version 1.1.0) with zero mismatches (-v 0). Reads maped to ribosomal RNAs (rRNA) were filtered out using the BOWTIE algorithm allowing three mismatches (-v 3). For counting mapping events to distinct RNA genes, the BOWTIE2 tool (*58*) was used with default settings. Reads were sorted to transfer RNAs (tRNAs), small nuclear/nucleolar RNAs (snoRNAs), messenger RNAs (mRNAs), all downloaded from the Ensembl database (*Botrytis cinerea* B05.10, ASM83294v1), as well as to retrotransposons, as annotated in a previous work (*40*). Reads counts were normalized on total Bc-sRNA reads per million (RPM).

### Sample preparation for mass spectrometry analysis

Protein samples were processed using the single-pot, solid-phase–enhanced sample preparation (SP3) protocol (*59*). Disulfide bonds were reduced by adding dithiothreitol (DTT) to a final concentration of 10 mM and incubating for 30 min at 56°C, followed by alkylation with 20 mM iodoacetamide for 30 min at 37°C in the dark. Samples were subsequently adjusted to 70% (v/v) acetonitrile (ACN), and 1 µl of carboxylate-modified magnetic beads (Sera-Mag Speed Beads, GE Healthcare) prepared as a 1:1 mixture of hydrophilic and hydrophobic beads in methanol/LC-MS grade water was added. Samples were mixed at 1,400 rpm for 18 min at room temperature to promote protein binding to the beads.

After binding, tubes were placed on a magnetic rack, and the supernatant was discarded. The beads were washed twice with 100% ACN and twice with 70% ethanol while kept on the magnet. For enzymatic digestion, beads were resuspended in 50 mM ammonium bicarbonate and incubated overnight at 37°C with sequencing-grade trypsin (Promega) at an enzyme-to-protein ratio of approximately 1:100 (w/w) while shaking at 1,400 rpm. Following digestion, peptides were rebound to the beads by adjusting the sample to 95% (v/v) ACN and shaking for 10 min at room temperature. The beads were collected on the magnet and washed twice with 100% ACN. Peptides were eluted with 2% DMSO in 1% formic acid, dried under vacuum, and stored at –20°C until LC–MS/MS analysis.

### Liquid Chromatography (UHPLC)

Peptide samples were reconstituted in loading solvent (0.1% formic acid in water) and analyzed by nano-UPLC using a Vanquish Neo UHPLC system (Thermo Fisher Scientific). A two-buffer solvent system was used, consisting of 0.1% formic acid in water as Buffer A and 0.1% formic acid in acetonitrile as Buffer B. For online desalting and concentration, each sample was first loaded onto a C18 trap cartridge (300 µm × 5 mm, 100 Å pore size, 5 µm particle, Thermo Fisher Scientific) and washed with Buffer A. The trapped peptides were then separated on a 25 cm analytical C18 reversed-phase column (75 µm inner diameter × 250 mm length, 130 Å pore size, 1.7 µm particle size; nanoEase BEH C18, Waters). Peptides were eluted with an 80-minute method employing a linear gradient from 2% to 30% Buffer B over 60 min, followed by high organic washes and re-equilibration.

### Mass Spectrometry (Orbitrap Exploris DDA)

Eluting peptides were analyzed on a hybrid quadrupole-Orbitrap mass spectrometer (Orbitrap Exploris 480, Thermo Fisher Scientific) operated in data-dependent acquisition (DDA) mode. The instrument was equipped with a nano-electrospray ionization source operating at a spray voltage of 1.8 kV. Full-scan MS1 spectra were acquired in the Orbitrap at a resolution of 60,000 (at m/z 200) over a mass range of m/z 350–1,400. The automatic gain control (AGC) target for MS1 was set to 3×10^6^ (normalized AGC 300%), with a maximum injection time of 25 msec. For MS2 fragmentation, the 20 most intense precursor ions (charge states 2–6) from each MS1 scan were selected in sequence for tandem MS. Precursors were isolated using a 2 m/z window in the quadrupole and fragmented by higher-energy collisional dissociation (HCD) with a normalized collision energy of 30%. The fragment ions were analyzed in the Orbitrap at a resolution of 15,000 (at m/z 200). The AGC target for MS2 was set to 5×10^4^ (normalized AGC 50%), with the maximum injection time automatically controlled by the instrument. The first mass for MS2 spectra was set to m/z 120. A dynamic exclusion of 30 sec was applied to avoid repeated sequencing of the same ion (±10 ppm exclusion window; isotopes were excluded; each precursor charge state was considered independently). An intensity threshold of 8×10^3^ was used to trigger MS2 scans, meaning only precursors exceeding this count were selected for fragmentation.

### Mass Spectrometry data analysis

Raw LC–MS/MS data were processed using Thermo Fisher Proteome Discoverer software (v3.0.0.757). Database searches were performed with the Sequest HT algorithm against the *Botrytis cinerea* reference proteome (UniProt Proteome ID: UP000001788, taxonomy ID: 332648; downloaded in February 2024). Carbamidomethylation of cysteine was set as a fixed modification. Methionine oxidation, N-terminal glutamine to pyro-glutamate conversion, and N-terminal protein acetylation were specified as variable modifications. Trypsin was defined as the protease, and up to two missed cleavages were allowed. Peptide length was restricted to between 6 and 144 amino acids. Search results were filtered to a 1% false discovery rate (FDR) at both the peptide and protein levels using a target-decoy approach.

### Proteomic analysis

Only proteins present across all three biological replicates with a minimum combined peptide sum of 3 were considered for analysis. Missing values were imputed with a constant value, defined as half of the lowest detected peptide number intensity across all samples (pseudocount = 0,5). Peptide counts were log₂-transformed, and differential abundance between conditions was assessed using the limma package (R/Bioconductor, version 3.21), which applies empirical Bayes moderation to stabilize variance estimates across proteins. The resulting p values were corrected using the Benjamini–Hochberg method to assess the false discovery rate (FDR). The fold-change was calculated on the average peptide count across 3 replicates. Proteins with FDR < 0.05 and a fold-change > 1 in crude compared to purified BcEVs were considered as “Crude BcEV-enriched proteins”. Gene ontology (GO)-term enrichment was performed on previously filtered EV proteins using g:Profiler (p-value threshold < 0.05). GPI-anchor containing proteins were predicted via PredGPI (*60*) and defined as putative adhesins. For comparison of identified EV proteins with previous publications, Protein IDs were acquired from supplemental datasets, considering only proteins classified by the authors as EV-associated. A summary of selected datasets can be found in Tab.S4. Where necessary, accession numbers not compatible with UniProt were transformed in protein identifiers through NCBI Batch Entrez. All Protein IDs were transformed in FASTA format using UniProt ID Mapping. De novo annotation of carbohydrate active enzymes (CAZymes) was performed using dbCAN3 (*61*), which utilizes 3 different models for protein annotation (DIAMOND, HMMER via CAZy and dbCAN-sub). Proteins were only classified as CAZymes if they were predicted by at least 2 out of the 3 models. Distinction between PCWDE, FCWDE and others was based on previous classifications (*62–64*).

### B. cinerea BcPME1-GFP reporter strain

The *BcPME1-GFP* strain was generated in the *B. cinerea* B05.10 background using the marker-free CRISPR/Cas9 knock-in strategy (*65*). Briefly, the endogenous *Bcpme1* locus (*BCIN_08g02970*) was targeted at the 3′ end of the coding region to introduce an in-frame C-terminal *GFP* transgene (*66*). The GFP repair template was amplified from plasmid pNAH-OGG (*67*) using primers carrying *BcPME1*-specific homology overhangs. The repair template was co-transformed with a Cas9/sgRNA complex and telomeric pTEL-Fen vector (*68*), which was used for fenhexamid selection. One sgRNA targeting the *BcPme1* locus was used for transformation. For transformation, 10 µg of the *BcPme1-GFP* repair template and 10 µg of pTEL-Fen vector were added to *B. cinerea* protoplasts at a concentration of 2×10⁷ protoplasts/ml. The Cas9/sgRNA complex was assembled using 2 µl sgRNA, 4 µl water, 3 µl cleavage buffer, and 6 µl Cas9 enzyme. Transformants were selected on medium containing 1 µg/ml fenhexamid. The genotype of the transformants was verified by diagnostic PCR using *BcPME1*-specific external primers and *GFP*-specific primers. Correct integration of GFP at the endogenous *BcPme1* locus was confirmed by sequencing. Primers used for strain generation and genotyping are listed in Tab.S6.

### Heterologous BcPME1 expression, purification and enzymatic activity

*BcPME1* was cloned into pEarlyGate104 plasmid harboring monomeric ultra-stable GFP (*69*) as an N-terminal fusion tag and transformed into the *Agrobacterium tumefaciens* strain LBA4404. *Agrobacteria* were cultivated overnight at 28°C on an LB-agar plate. Colonies were resuspended in 1 ml of ¼ MS pH 5.7, 1% (w/v) sucrose, 100 µM acetosyringone, 0.005% (v/v) Silwet L-77. Cell suspensions were adjusted to an OD600 nm of 0.5 and ten 4 - 6 weeks old tobacco plants were leaf-infiltrated. Plants were cultivated in dark for 24°C at 75% humidity followed by 2 to 3 days at long day conditions at 170 µE light intensity using Philips LED tubes HO 18.5 W 2800 lm 840K. Leaves were harvested and frozen in liquid nitrogen. Frozen plant material was ground and 1.2 ml lysis buffer (50 m Tris-HCl pH 7.5, 150 mM NaCl, 2.5 mM MgCl2, 1 mM DTT, 0.5% NP-40) per 0.25 g plant material was added. One tablet complete protease inhibitor (Roche) was added and lysis was allowed to continue for 30 min on ice with inverting the tube every 5 min. Lysed plant material was filtered through two cell sieves (70 µm mesh size). For BcPME1 isolation, 100 µg of GST-tagged anti-GFP nanobody (*70*) were immobilized on 30 µl glutathione high-capacity magnetic agarose beads (Sigma-Aldrich) prior adding to plant lysate. Lysate and coupled magnetic beads were incubated for 2 hours at 4°C with gentle shaking. After incubation, immobilized GFP-BcPME1 was washed thrice with 1 ml of ice-cold lysis buffer and either eluted in 50 µl of lysis buffer supplemented with 10 mM glutathione or remained bead bound. Isolated BcPME1 was snap frozen in liquid nitrogen until further use.

BcPME1 activity was measured using a modified assay according to Tsuji *et al*. (*71*) and Grsic-Rausch *et al*. (*72*). In brief, pectin from citrus peel at a final concentration of 0.44% (w/v) was added to 50 µl of a reaction mixture consisting of 8.8 mU alcohol oxidase from *Candida boidinii* (Sigma-Aldrich), 39 µM AmpliFlu Red (Sigma-Aldrich) and 0.2 U/ml horse radish peroxidase (Sigma-Aldrich) in 1x PBS at pH 7.0. Immobilized BcPME1 was added to the reaction mixture and fluorescence was measured every 30 sec for 30 min at room temperature on a Tecan Spark with emission wavelength set to 570 nm (bandwidth 5 nm) and excitation wavelength set to 590 nm (bandwidth 5 nm).

### Plots and statistical analysis

Computational analyses and data plotting was done in R studio (version: 2025.09.0+387). Plots and figures were formatted in Affinity Designer 2 (version: 2.6.2). Chat GPT (Version 4) and Claude Sonnet 5 assisted in text correction and code troubleshooting.

### LLM

Chat GPT-4.1 nano was used for manuscript text polishing. Authors have carefully reviewed the text upon polishing for its accuracy.

---

**Figure S1:**
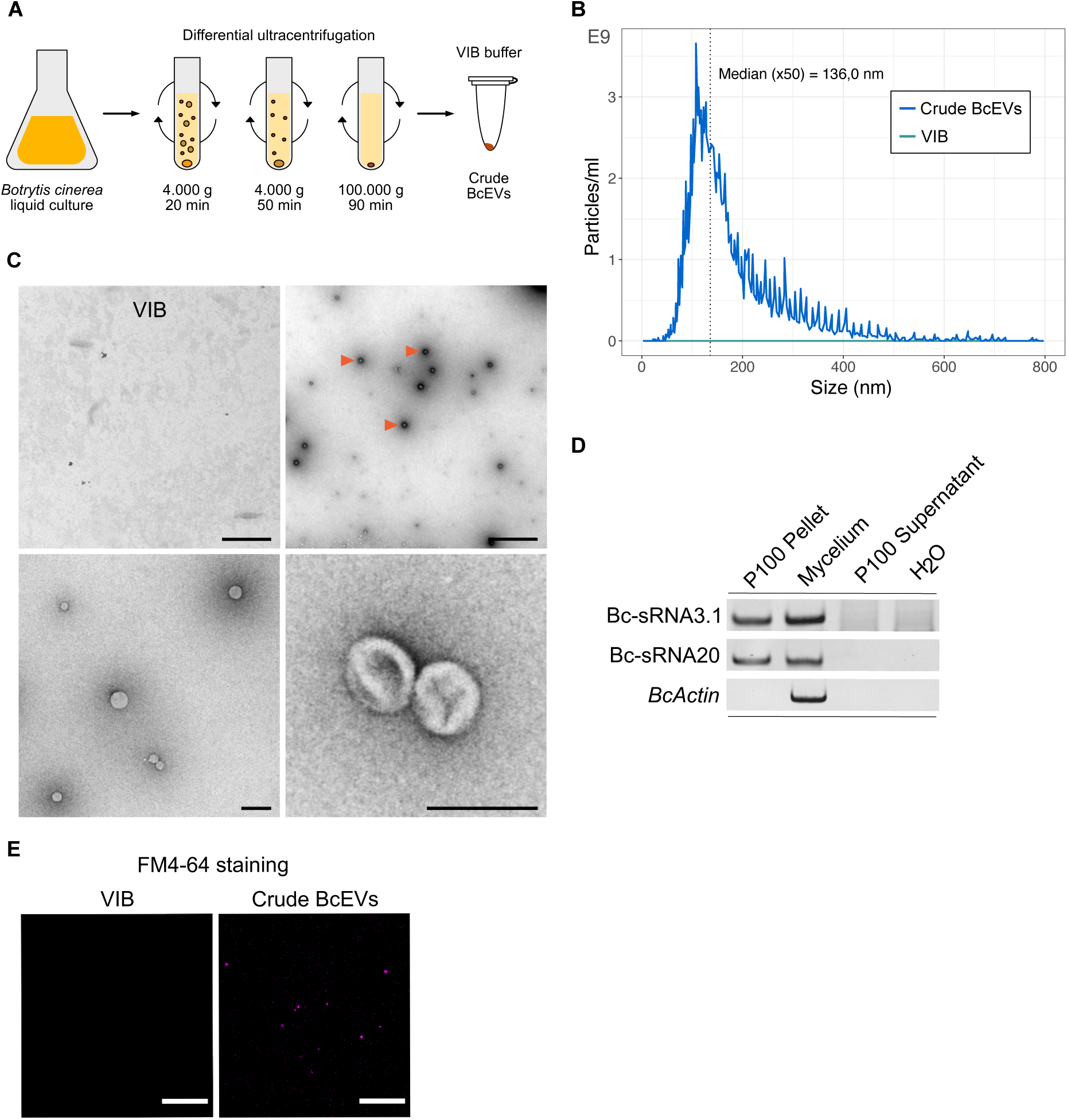
Crude BcEV isolation from *B. cinerea* liquid culture. A) Workflow of crude BcEV isolation from fungal liquid culture. B) Representative NTA blot of crude BcEVs. C) Representative TEM images of crude BcEVs. The red arrows indicate BcEV particles. The scale bars indicate 500 nm in the top and 200 nm in the bottom row. D) Stem-loop RT-PCR of Bc-sRNA3.1 (*73*) and Bc-sRNA20 (*40*) in crude BcEVs (P100 pellet). E) FM4-64 staining of crude BcEVs. The scale bars indicate 5 µm.

**Figure S2:**
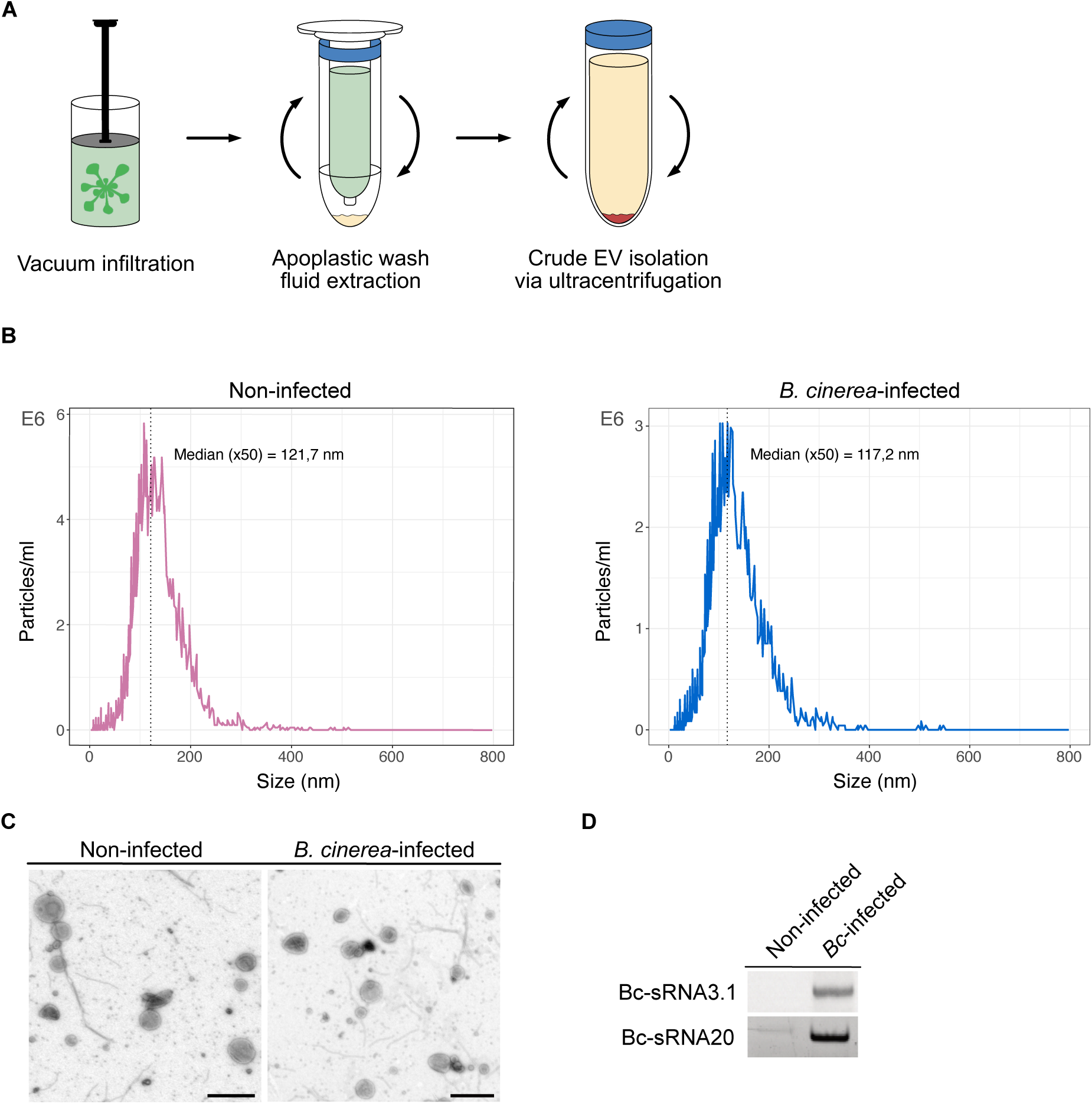
Crude BcEV isolation from *A. thaliana* apoplastic wash fluids. A) Workflow to collect crude BcEV from apoplastic wash fluids (AWF) of *B. cinerea*-infected *A. thaliana*. B) NTA of AWF-isolated EVs from non-infected and *B. cinerea*-infected *A. thaliana*. C) TEM images of AWF-isolated EVs from non-infected and *B. cinerea*-infected *A. thaliana*. Scale bars indicate 500 nm. D) Stem-loop RT-PCR of Bc-sRNA3.1 and Bc-sRNA20 in AWF-isolated EVs from non-infected and *B. cinerea*-infected *A. thaliana*.

**Figure S3:**
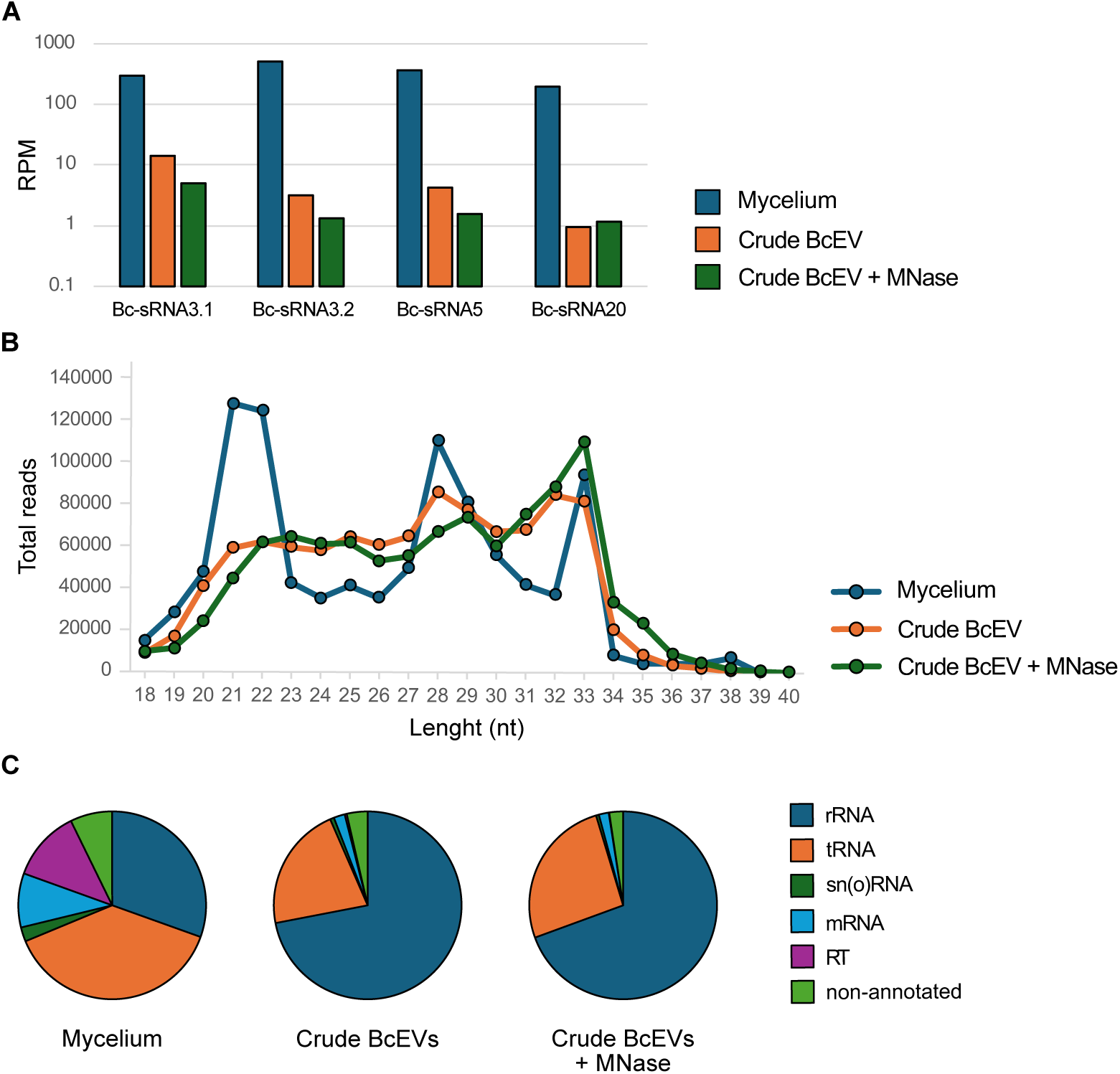
Small RNA sequencing of crude BcEVs. A) Reads per million (RPM) of Bc-sRNA3.1, Bc-sRNA3.2, Bc-sRNA5 (*73*) and Bc-sRNA20 in mycelium, or crude BcEV samples with or without MNaseI treatment, based on Illumina short read sequencing. B) Small RNA size profiles of Illumina sequencing. C) Distribution of sRNA origins, based on Illumina sequencing.

**Figure S4:**
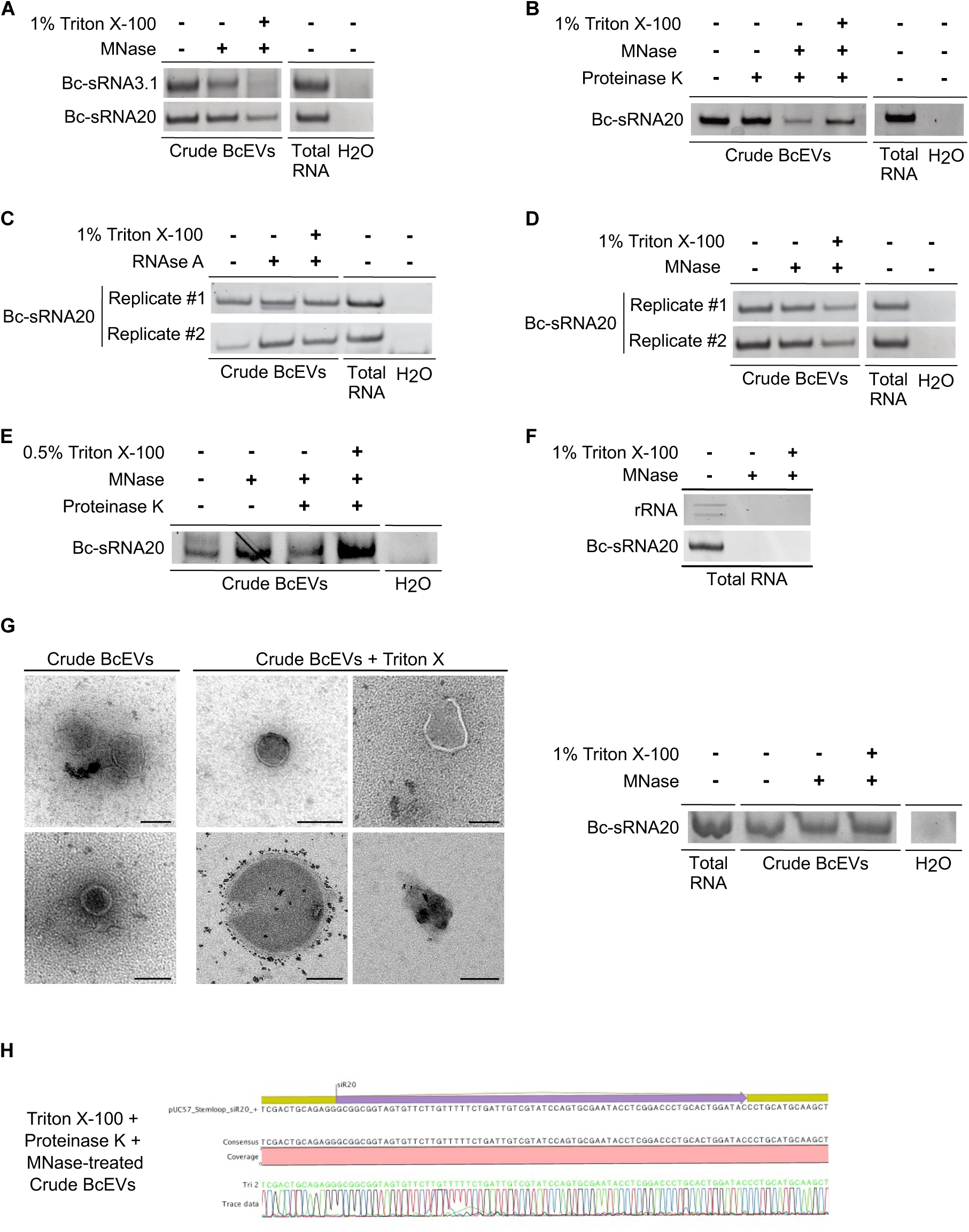
Small RNA protection assays with crude BcEVs. A) Stem-loop RT-PCR of Bc-sRNA3.1 and Bc-sRNA20 in crude BcEV samples upon treatment with MNaseI and 1% Triton X-100. Total RNA was used as positive and water (H2O) as negative control. B) Bc-sRNA20 was detected upon sequential treatments with Proteinase K and MNaseI following 1% Triton X-100 incubation. C, D) Two biological replicates of stem-loop RT-PCRs detecting Bc-sRNA20 in crude BcEV samples upon MNaseI or RNAse A treatment with or without 1% Triton X-100 incubation. E) Bc-sRNA20 was detected upon sequential treatments with Proteinase K and MNaseI following 0.5% Triton X-100 incubation. F) Stem-loop RT-PCR of Bc-sRNA20 with total RNA isolated from *B. cinerea* mycelium upon treatment with MNaseI and 1% Triton X-100 indicating that MNaseI degraded unprotected ribosomal RNA (rRNA) and Bc-sRNA20. G) Representative TEM images of crude BcEVs after 1% Triton X treatment indicated partially disrupted BcEVs. Bc-sRNA20 was detected upon treatment with MNaseI following 1% Triton X-100 incubation in the same samples used for TEM imaging. Scale bars represent 100 nm. H) Sanger-sequencing result of Bc-sRNA20 cloned from crude BcEVs upon MNaseI, proteinase K and 1% Triton X-100 treatment.

**Figure S5:**
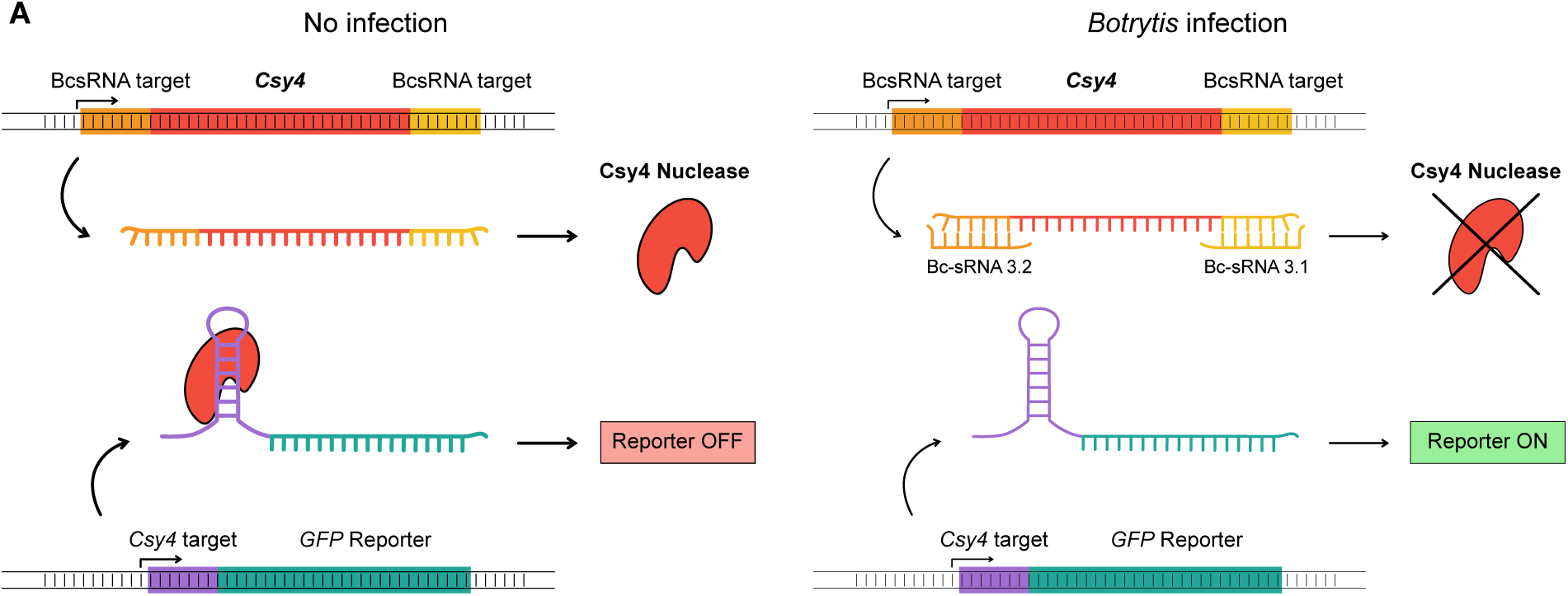
GFP switch-on cross-kingdom RNAi reporter construct. Schematic view of the GFP switch-on cross-kingdom RNAi reporter construct stably expressed in *A. thaliana* reporter plants.

**Figure S6:**
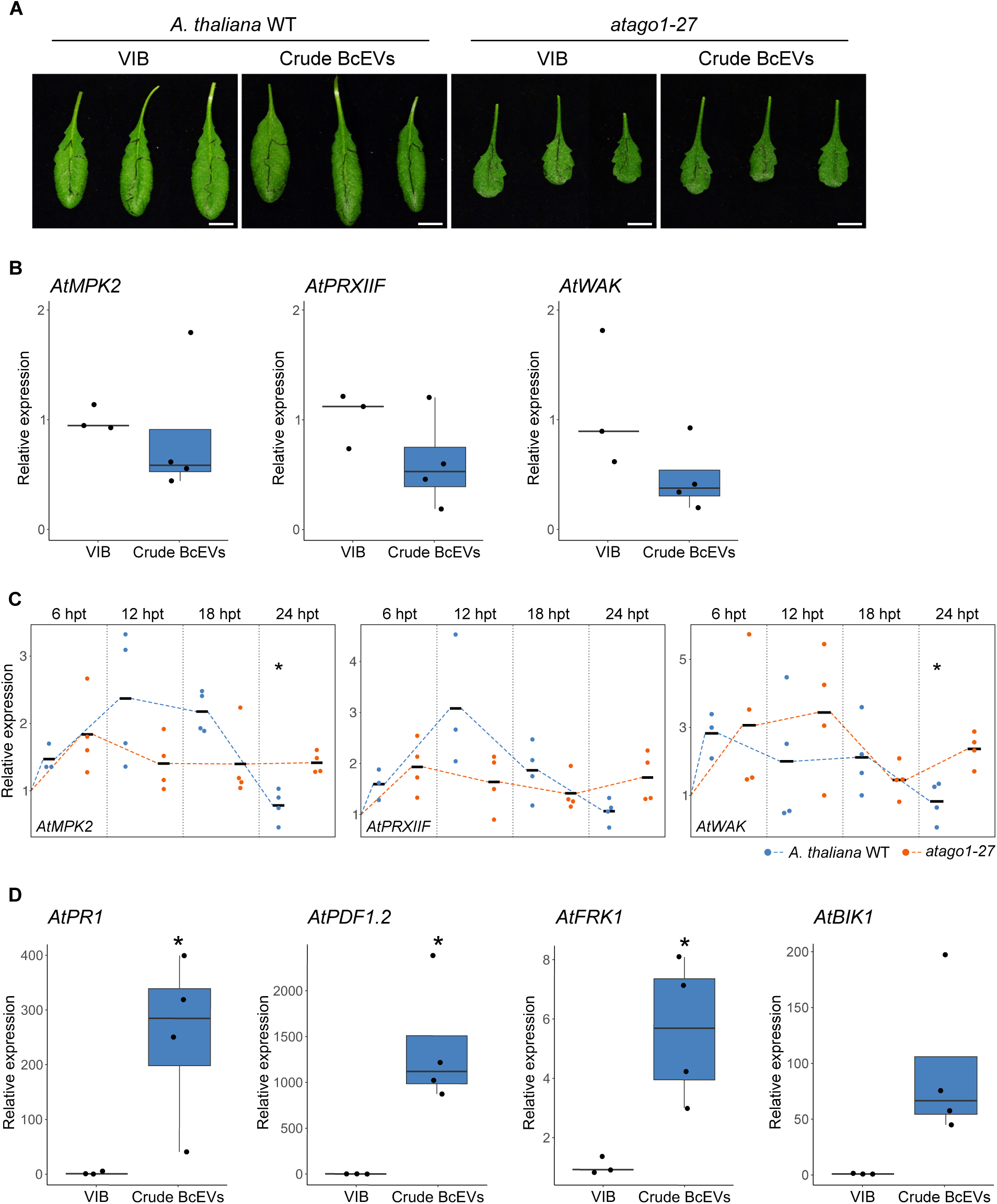
Target gene suppression by crude BcEVs. A) *A. thaliana* WT and *atago1-27* mutant leaves infiltrated with crude BcEVs. Scale bars indicate 1 cm. B) qRT-PCR analysis of the Bc-sRNA target genes in *A. thaliana AtWAK*, *AtMPK2*, *AtPRXIIF* upon infiltration of crude BcEVs. C) Time course qRT-PCR analysis of the Bc-sRNA target genes in *A. thaliana* WT and *atago1-27* mutant plants. Data points represent four biological replicates, with bars representing the mean value and significance assessed by pairwise student’s t-test with p<0.05 indicated by asterisks. D) Gene expression analysis of immune-related genes in *A. thaliana* WT upon treatment with crude BcEVs at 18 hpi. The plant *AtCDKA* gene was used as reference in B), C), and D). Data points represent biological replicates. Asterisks indicate significant differences using a pairwise t-test with p< 0.05.

**Figure S7:**
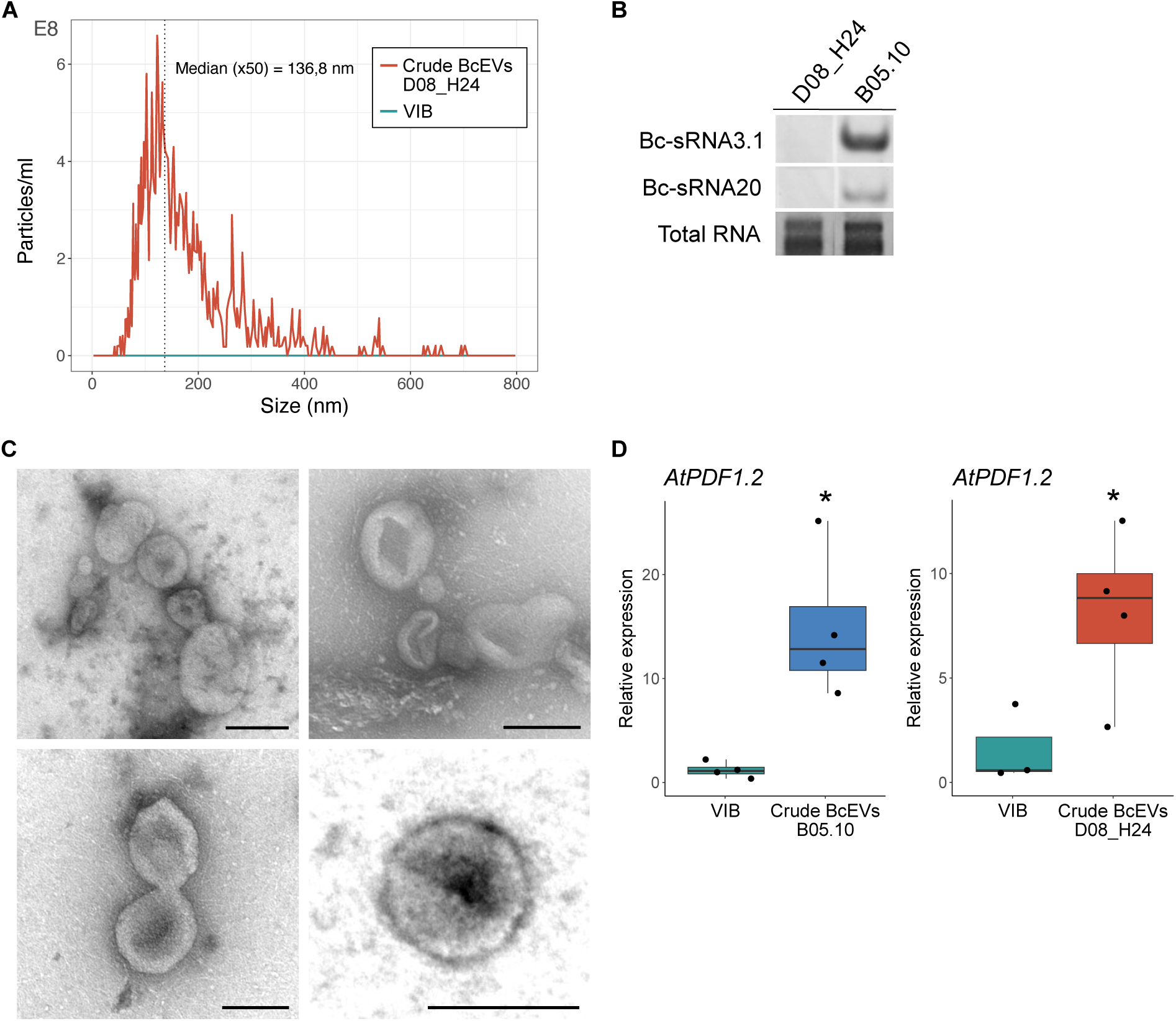
Crude BcEV isolation of the strain *B. cinerea* D08_H24. A) Representative NTA plot of crude BcEVs isolated from the *B. cinerea* strain D08_H24. NTA measurements were repeated in two independent experiments with comparable results. B) Stem-loop RT-PCR of Bc-sRNA3.1 and Bc-sRNA20 in *B. cinerea* B05.10 and D08_H24 mycelium. C) Representative TEM images of crude BcEVs of D08_H24. Scale bars indicate 100 nm. D) Gene expression analysis of the immune-related gene *AtPDF1.2* in *A. thaliana* upon treatment with crude BcEVs of B05.10 or D08_H24 at 24 hpi by qRT-PCR. The *AtCDKA* gene was used as reference. Data points represent four biological replicates. Asterisks indicate significant differences using a pairwise t-test with p< 0.05.

**Figure S8:**
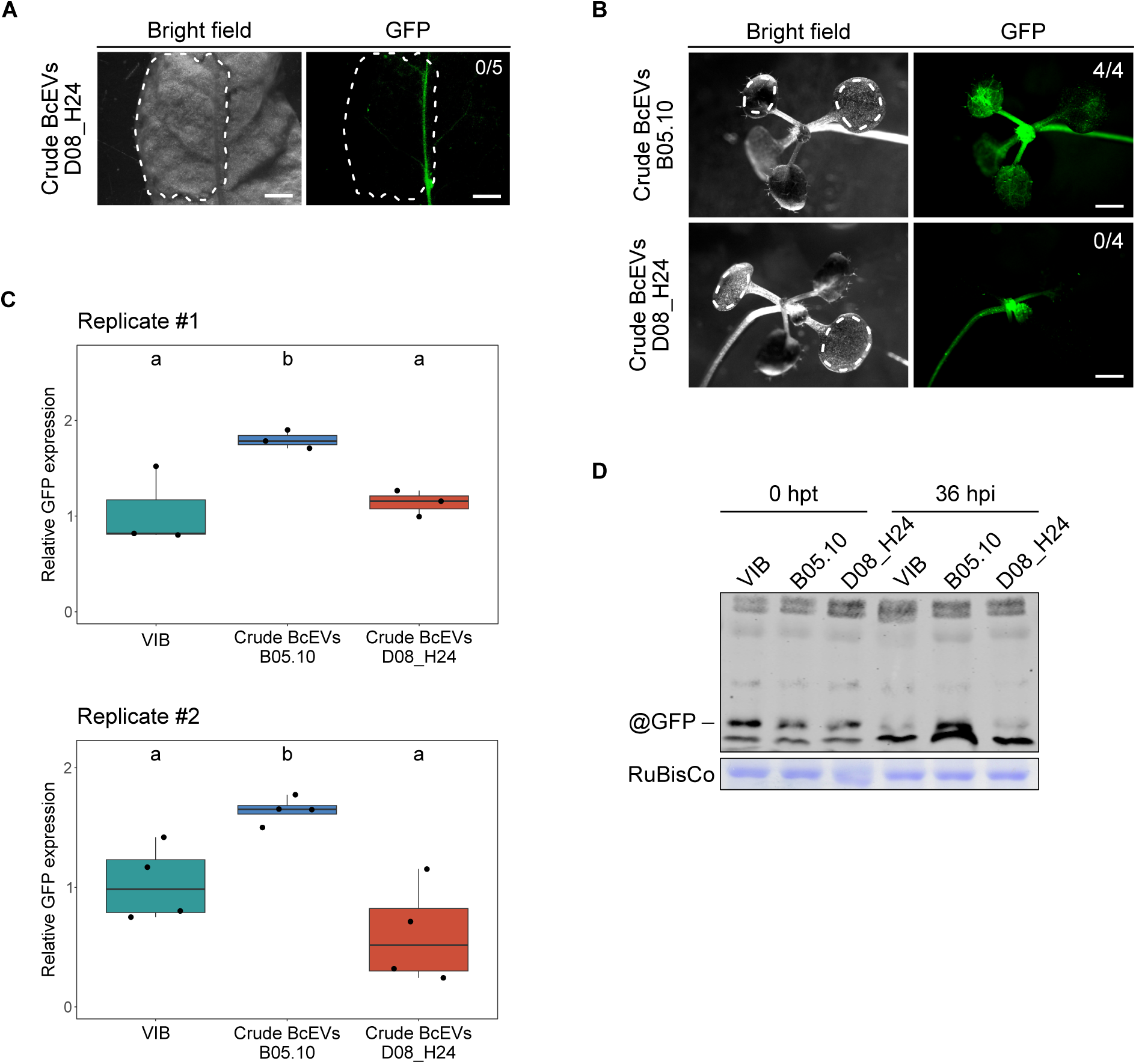
GFP reporter plants treated with crude BcEVs of *B. cinerea* B05.10 and D08_H24 strains. A) Leaf infiltration of crude BcEVs of *B. cinerea* D08_H24 into GFP reporter plants. Image taken at 24 hpi. Dotted lines indicate area of infiltration. B) Leaf drop assay of crude BcEVs of *B. cinerea* B05.10 or D08_H24 strains using *A. thaliana* GFP reporter seedlings, without detaching the leaves prior to treatment. Images taken at 36 hpi. C) *GFP* gene expression in *A. thaliana* reporter plants treated with crude BcEVs of *B. cinerea* B05.10 or D08_H24 strains at 24 hpi by qRT-PCR. The *AtCDKA* gene was used as reference. Data points represent biological replicates. Asterisks indicate significant differences using a two-way ANOVA test with Tukey HSD p< 0.05. D) Immunoblot analysis of GFP using total protein extracts of GFP reporter plants infiltrated with VIB or crude BcEVs of B05.10 or D08_H24 strains. RuBisCo was used as a loading control.

**Figure S9:**
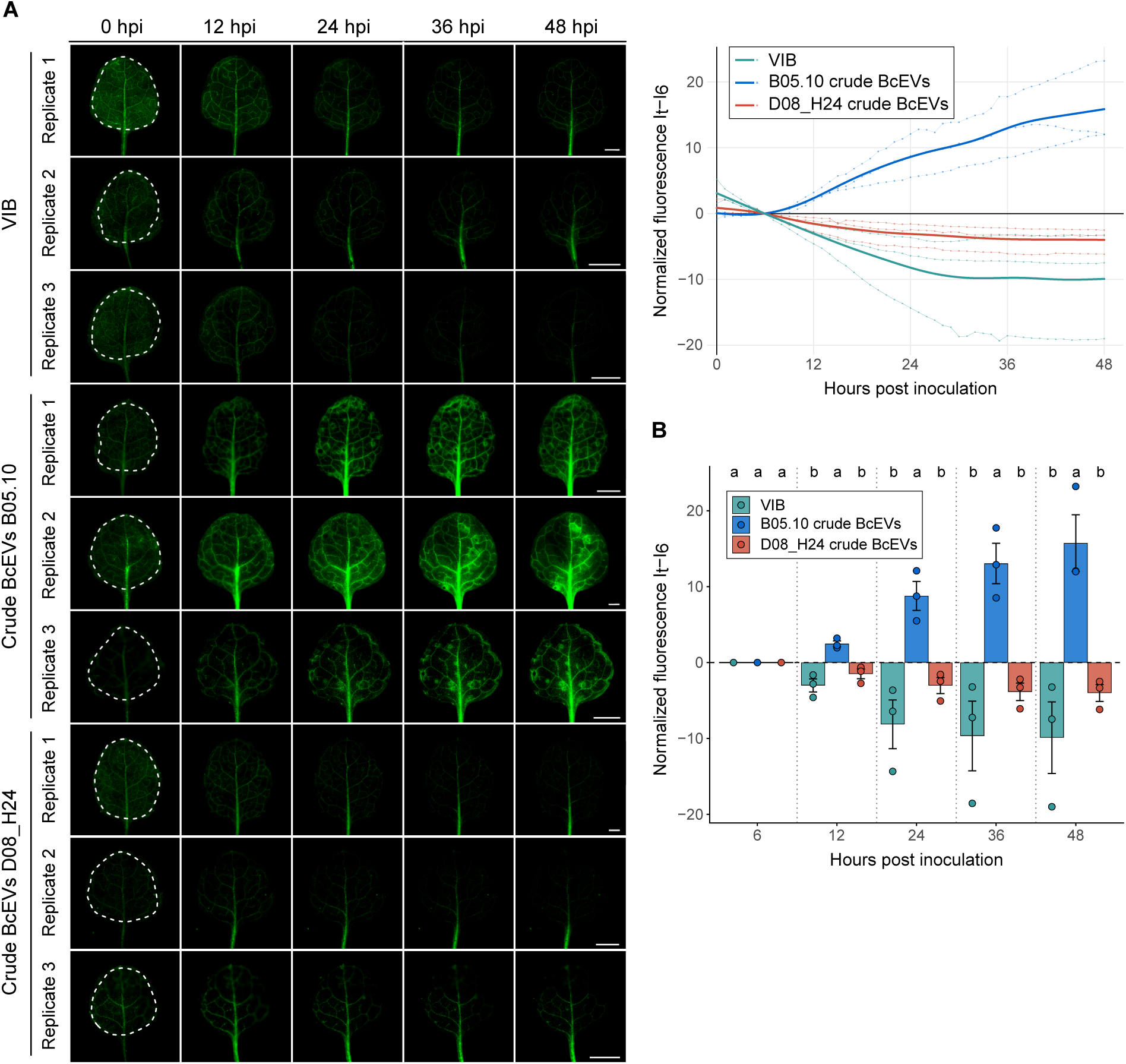
Time-course fluorescence microscopy of GFP reporter plants treated with crude BcEVs of *B. cinerea* B05.10 and D08_H24 strains. A) 50 µl crude BcEVs diluted in 40 µl instead of 200µl VIB (1,1×10^13^ particles/ml) comparing the *B. cinerea* strains B05.10 or D08_H24 were dropped on the entire surface of GFP reporter leaves and covered with a glass slide. Time-course fluorescence microscopy was carried out from 0 – 48 hpi. Scale bars in the images represent 1 mm. GFP signals were imaged and quantified each 60 min. Normalized GFP fluorescence intensity (It – I6) was plotted to evaluate GFP activation. Dots connected by thin lines represent individual biological replicates (n = 3); bold lines represent mean values. Replicates of time-course fluorescence microscopy of GFP reporter plants treated with crude BcEVs of *B. cinerea* B05.10 or D08_H24 strains. Dotted lines indicate area of treatment. Scale bars represent 2 mm. Normalized GFP fluorescence intensity (It – I6) at selected time points (6, 12, 24, and 36 hpi), corresponding to the data shown in Fig.S9A. B) Bar graphs represent mean ± SEM of three biological replicates. Statistical significance was determined by pairwise ANOVA followed by Tukey’s HSD post hoc test at each time point; treatments not sharing the same letter are significantly different (p < 0.05).

**Figure S10:**
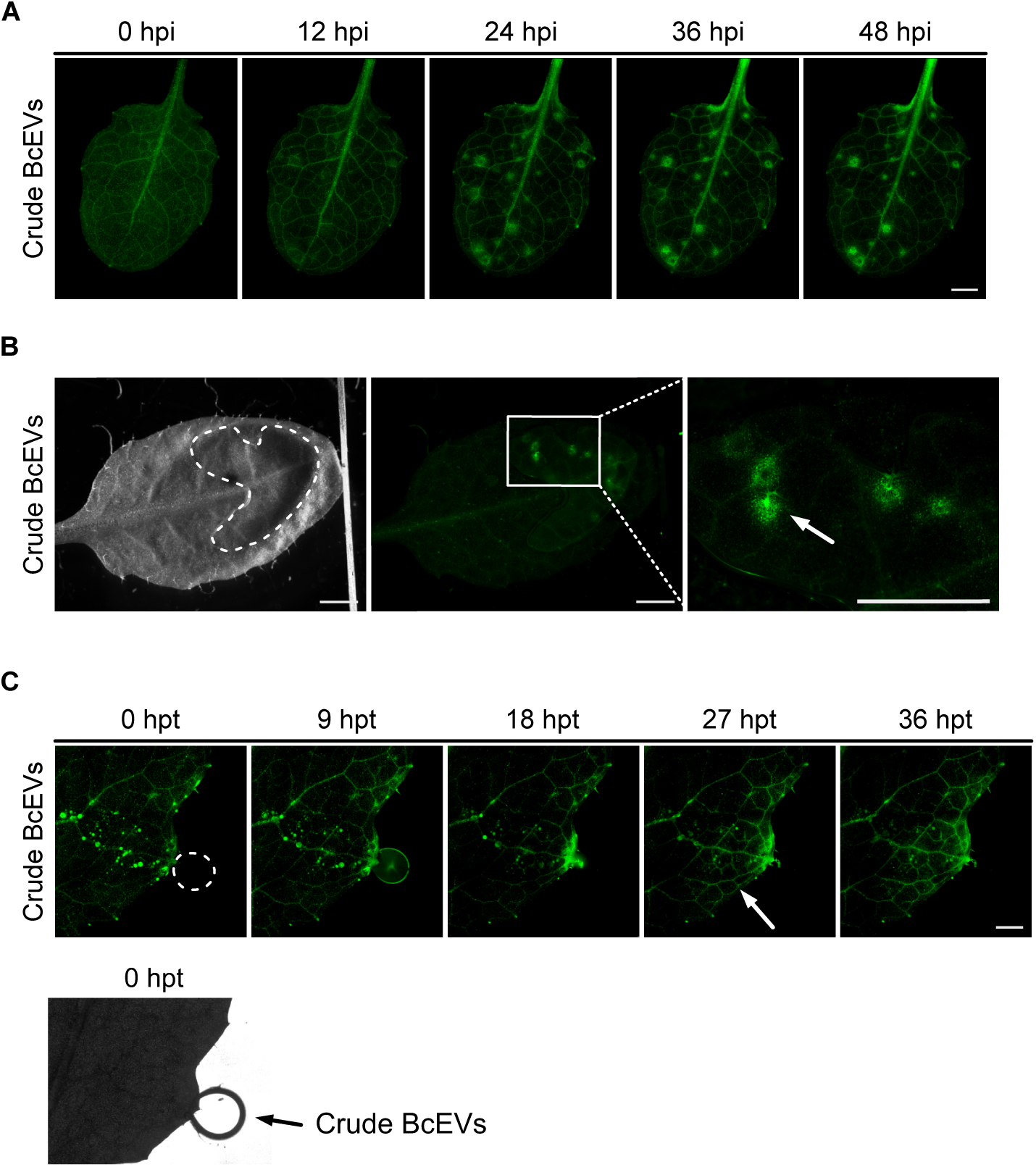
Replicates of GFP reporter plants treated with crude BcEVs. A) Time-course fluorescence microscopy of GFP reporter plants after drop inoculation with 10 µl crude BcEVs (1,1×10^13^ particles/ml) on the entire leaf surface. Scale bar indicates 1 mm. B) Drop inoculation of GFP reporter plants with 5 µl crude BcEVs. Bright field and fluorescence image were taken with an epifluorescence stereomicroscope with a GFP filter. Arrow indicates GFP activity around trichomes. C) Time-course fluorescence microscopy of GFP reporter plants after drop inoculation with 10 µl crude BcEVs on leaf edge in hydathode proximity (dotted circle). Arrow indicates GFP spreading at leaf veins. Fluorescence images were taken with a Thunder Imaging platform. Scale bar indicates 100 µm. Fig.S10A is equivalent to the video S1 and Fig.S10C to the video S2.

**Figure S11:**
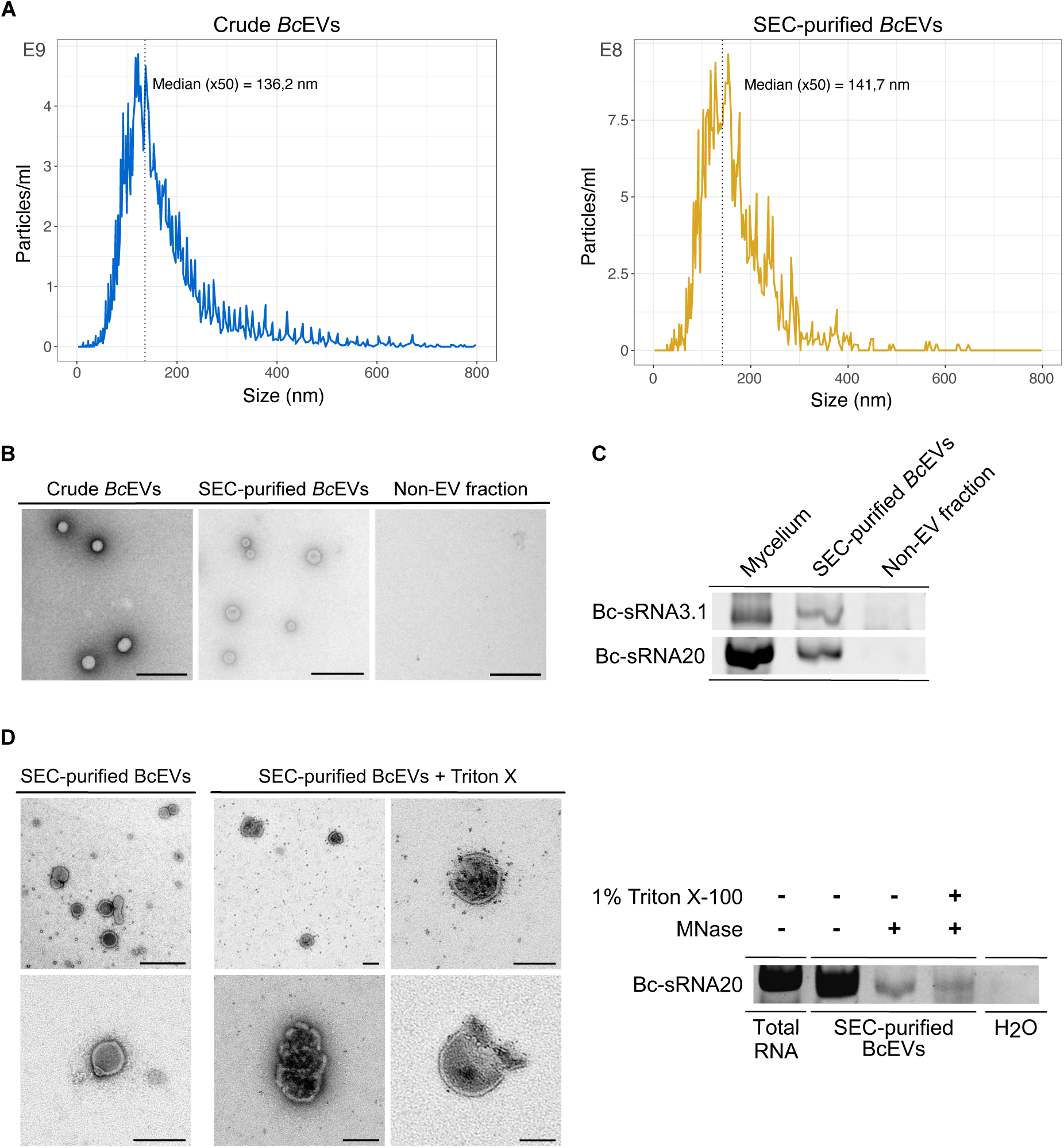
Analysis of SEC-purified BcEVs. A) NTA plots of crude BcEVs and SEC-purified BcEVs of the same sample isolated from *B. cinerea* B05.10 liquid culture. B) Representative TEM images of crude BcEVs and SEC-purified BcEVs of the same sample. Scale bars represent 500 nm. C) Stem-loop RT-PCR of Bc-sRNA3.1 and Bc-sRNA20 in crude BcEVs and SEC-purified BcEVs of the same sample. D) Representative TEM images of SEC-purified BcEVs after 1% Triton X treatment show partially disrupted or collapsed BcEVs. Bc-sRNA20 was detected upon treatment with MNaseI following 1% Triton X-100 incubation in the same samples used for TEM imaging. Scale bars represent 500 nm for top left image and 100 nm for all other images.

**Figure S12:**
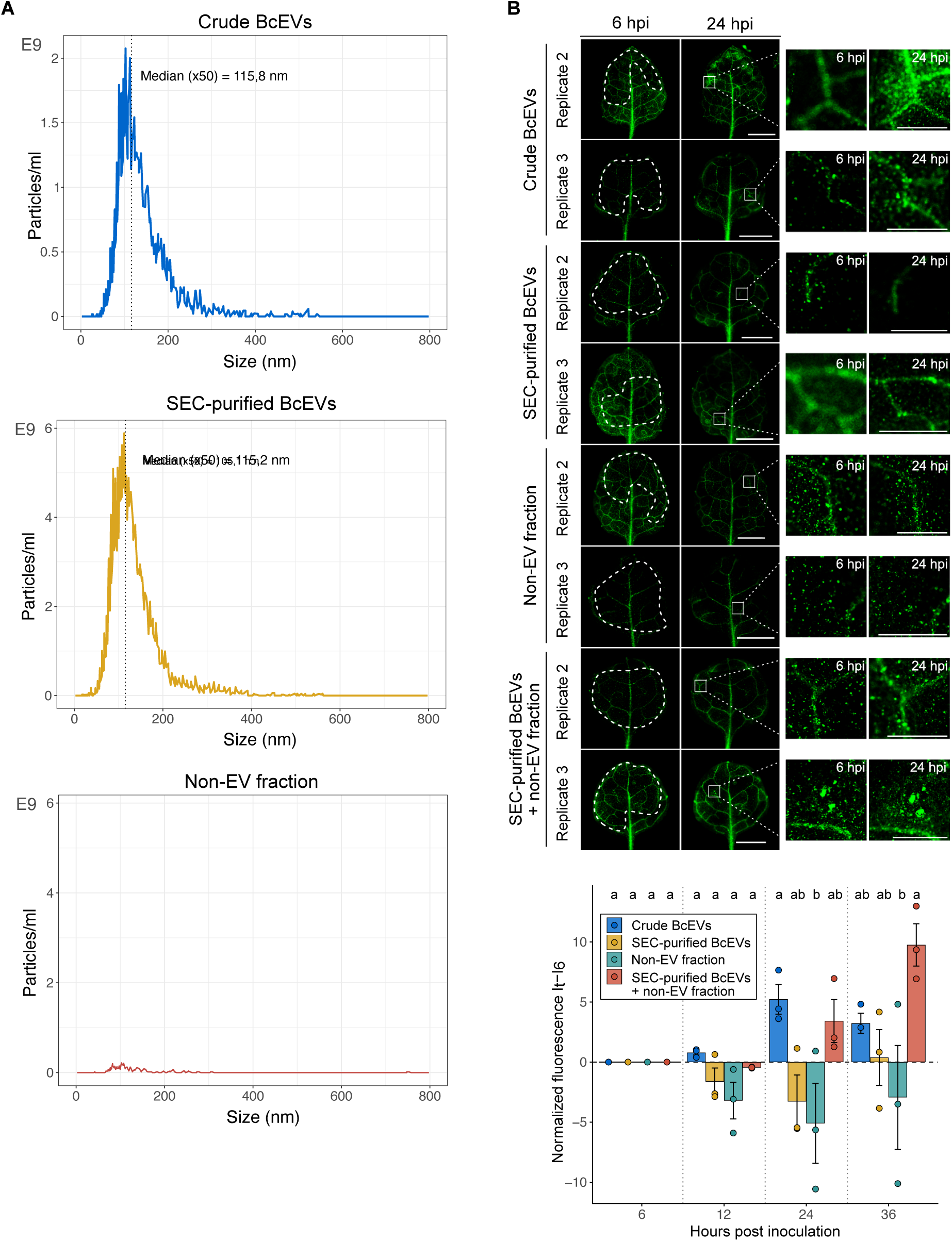
Replicates of GFP reporter plants treated with crude BcEVs or SEC-purified BcEVs. A) NTA measurements of crude and SEC-purified BcEVs as well as the non-EV fraction. B) Replicates of GFP reporter plants treated with crude BcEVs or SEC-purified BcEVs as well as non-EV fraction, according to Fig.1F. Dashed white lines outline the area of BcEV treatment; the white square at 24 hpi marks a representative site of GFP activation, shown magnified in the adjacent panels. Normalized GFP fluorescence intensity (It – I6) at selected time points (6, 12, 24, and 36 hpi), corresponding to the data shown in Fig.1F and Fig.S12B. Bar graphs represent mean ± SEM of three biological replicates. Statistical significance was determined by pairwise ANOVA followed by Tukey’s HSD post hoc test at each time point; treatments not sharing the same letter are significantly different (p < 0.05).

**Figure S13:**
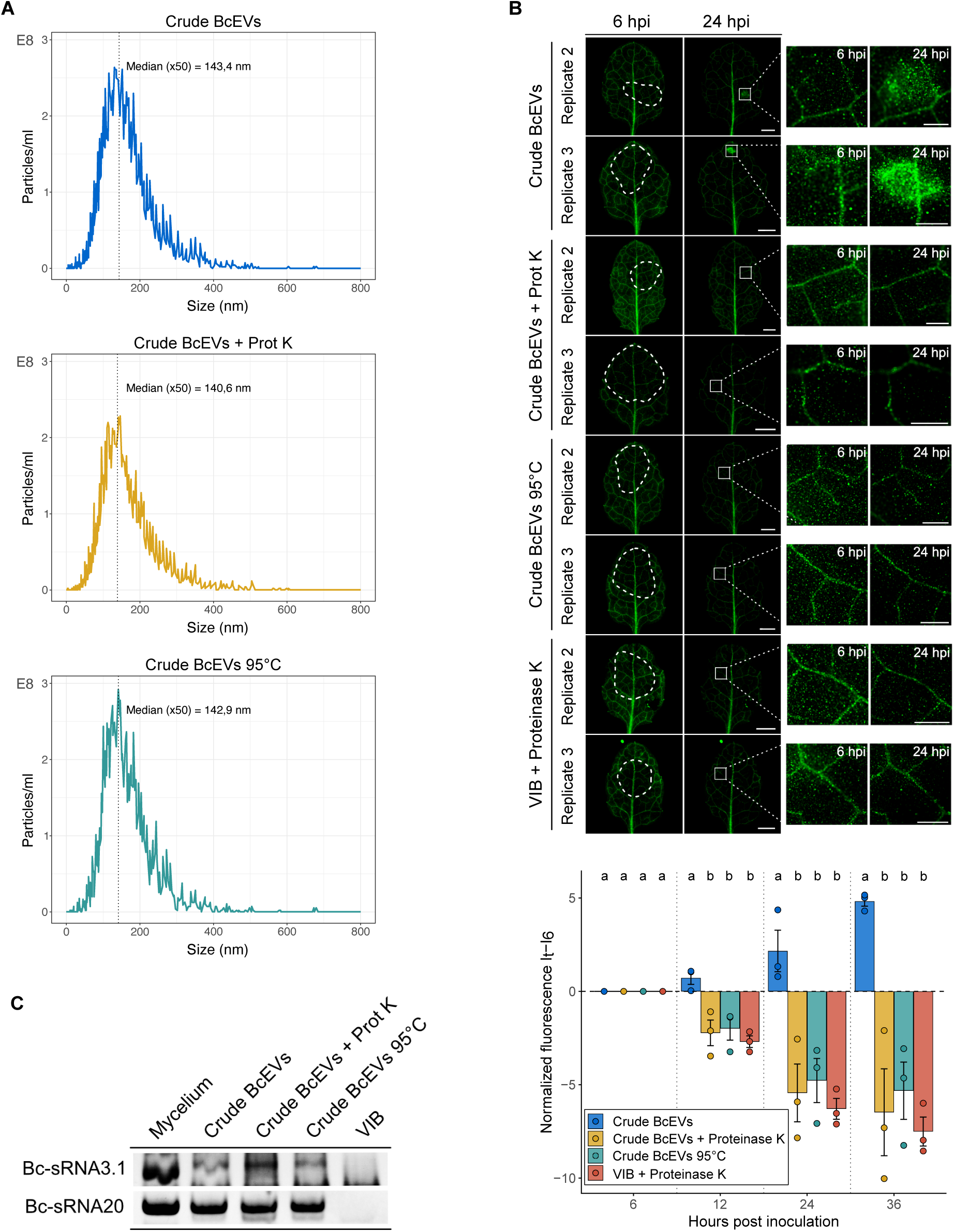
Analysis of heat-treated or proteinase K-treated crude BcEVs. A) NTA plots of crude BcEVs upon heat-treatment or incubation with Proteinase K. B) Replicates of GFP reporter plants treated with crude BcEVs upon heat-treatment or incubation with Proteinase K, according to Fig.2B. Dashed white lines outline the area of BcEV treatment; the white square at 24 hpi marks a representative site of GFP activation, shown magnified in the adjacent panels. Scale bars represent 2 mm. Normalized GFP fluorescence intensity (It – I6) at selected time points (6, 12, 24, and 36 hpi), corresponding to the data shown in Fig.1F and Fig.S13B. Bar graphs represent mean ± SEM of three biological replicates. Statistical significance was determined by pairwise ANOVA followed by Tukey’s HSD post hoc test at each time point; treatments not sharing the same letter are significantly different (p < 0.05). C) Stem-loop RT-PCR of Bc-sRNA3.1 and Bc-sRNA20 with crude BcEVs upon heat-treatment or incubation with Proteinase K.

**Figure S14:**
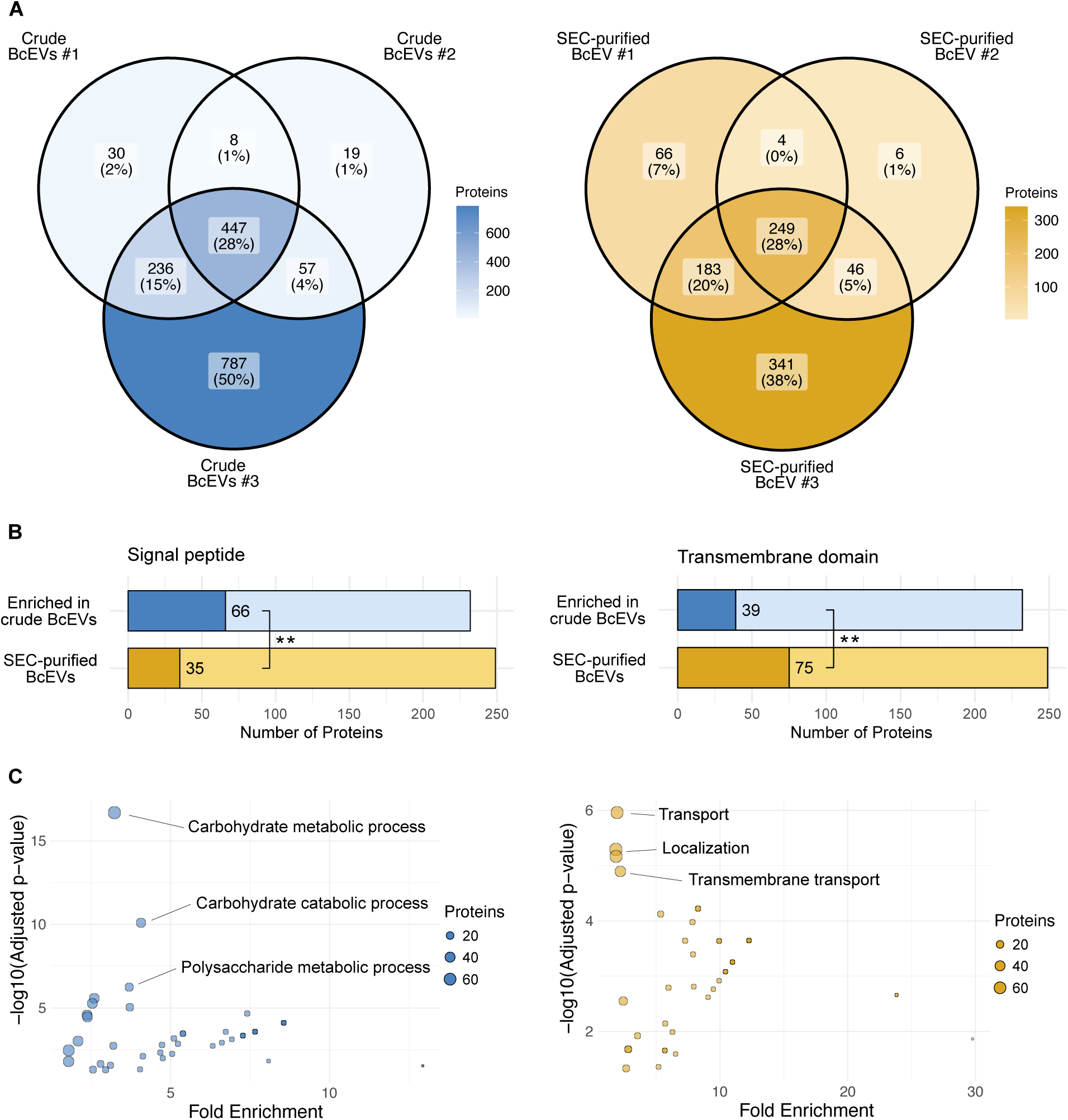
Protein analysis of crude BcEV versus SEC-purified BcEV samples. a) Venn diagrams indicating numbers of overlapping and distinct protein hits in three biological replicates of crude BcEV and SEC-purified BcEV samples. b) Counts of protein hits with predicted signal peptides (SP) and transmembrane domains in crude BcEV and SEC-purified BcEV samples. c) Enrichment analysis of protein hits identified in crude BcEV and SEC-purified BcEV samples.

**Figure S15:**
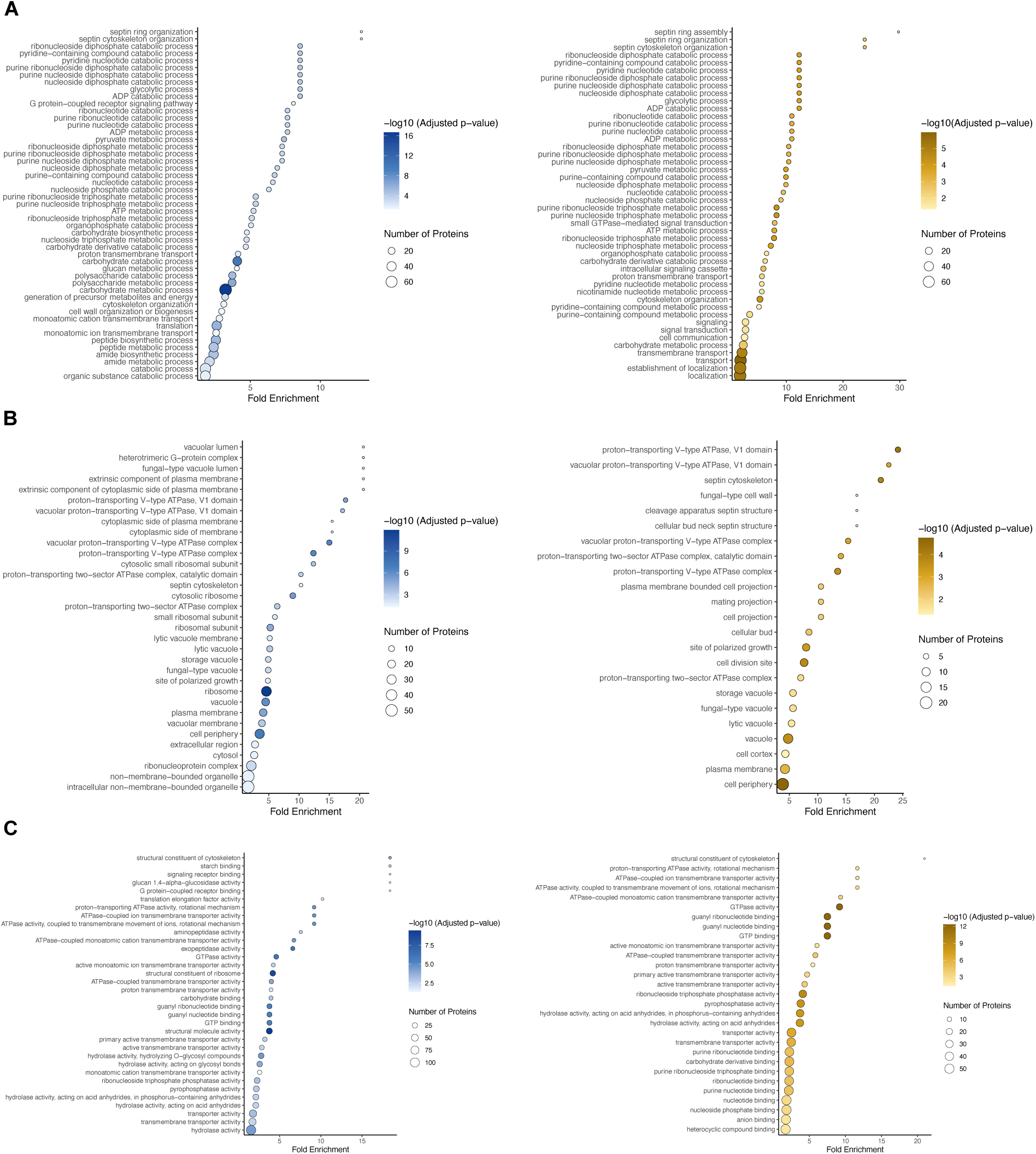
GO enrichment analysis of crude BcEV proteins. A) GO enrichment analysis of crude and SEC-purified BcEV protein data according to biological processes. B) GO enrichment analysis of crude and SEC-purified BcEV protein data according to cellular compartment. C) GO enrichment analysis of crude and SEC-purified BcEV protein data according to molecular function.

**Figure S16:**
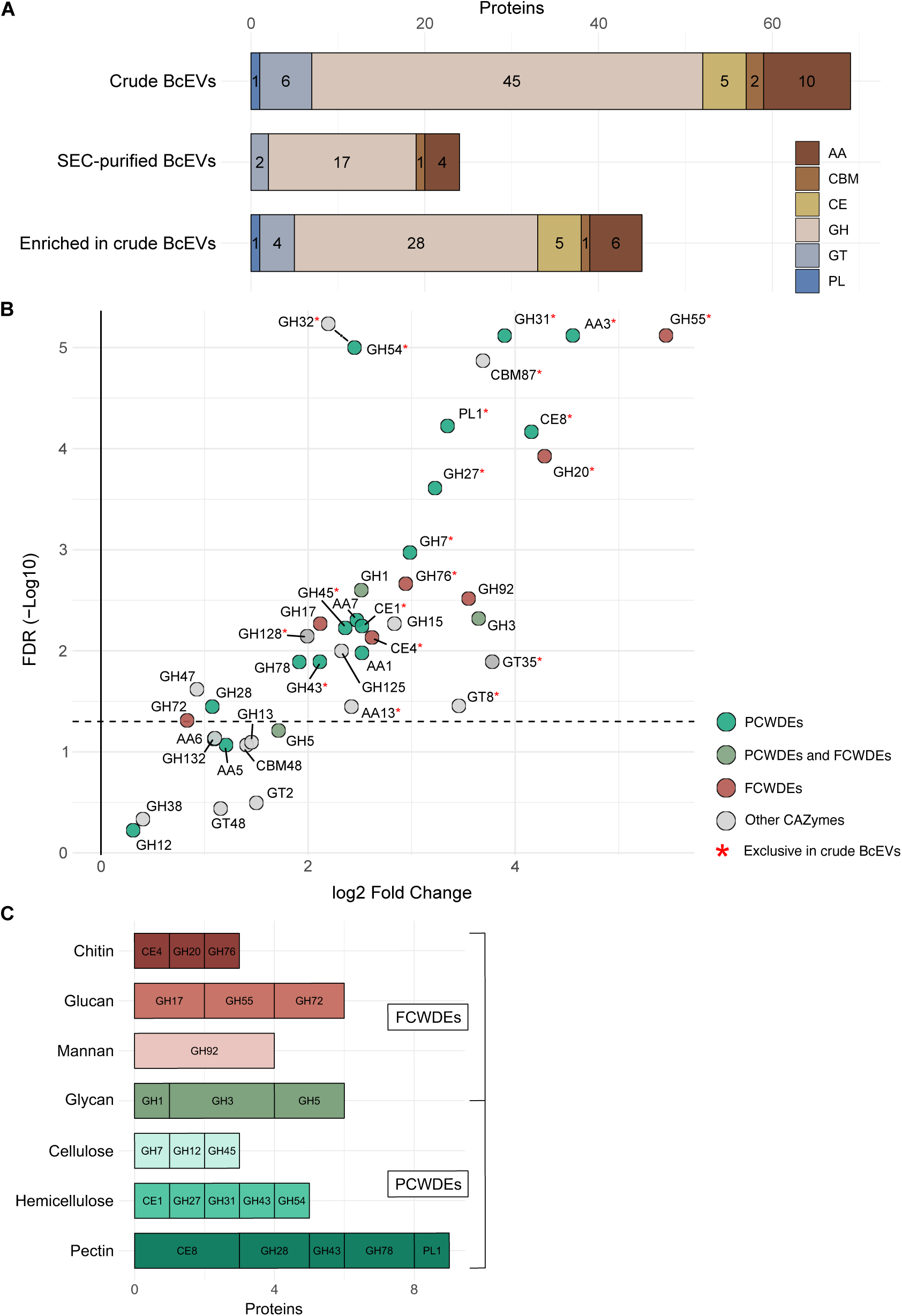
CAZymes identified in BcEV samples. A) CAZyme classes detected in crude BcEV, SEC-purified BcEV, and proteins enriched exclusively in crude BcEV samples. CAZyme classes include auxiliary activity enzymes (AA), carbohydrate-binding modules (CBM), carbohydrate esterases (CE), glycoside hydrolases (GH), glycosyltransferases (GT), and polysaccharide lyases (PL). B) Volcano plot of CAZyme families identified in crude versus SEC-purified BcEV samples. Peptide counts for proteins within the same CAZyme family were summed for each of the three biological replicates. The p-values were calculated based on a Benjamini-Hochberg corrected pairwise t-test. Sub-families detected exclusively in crude BcEVs are marked with a red asterisk. C) CAZyme sub-families present in crude BcEVs, with their constituent proteins categorized as plant cell wall–degrading enzymes (PCWDEs) or fungal cell wall–degrading enzymes (FCWDEs), and the corresponding carbohydrate substrates they act on in the plant or fungal cell wall.

**Figure S17:**
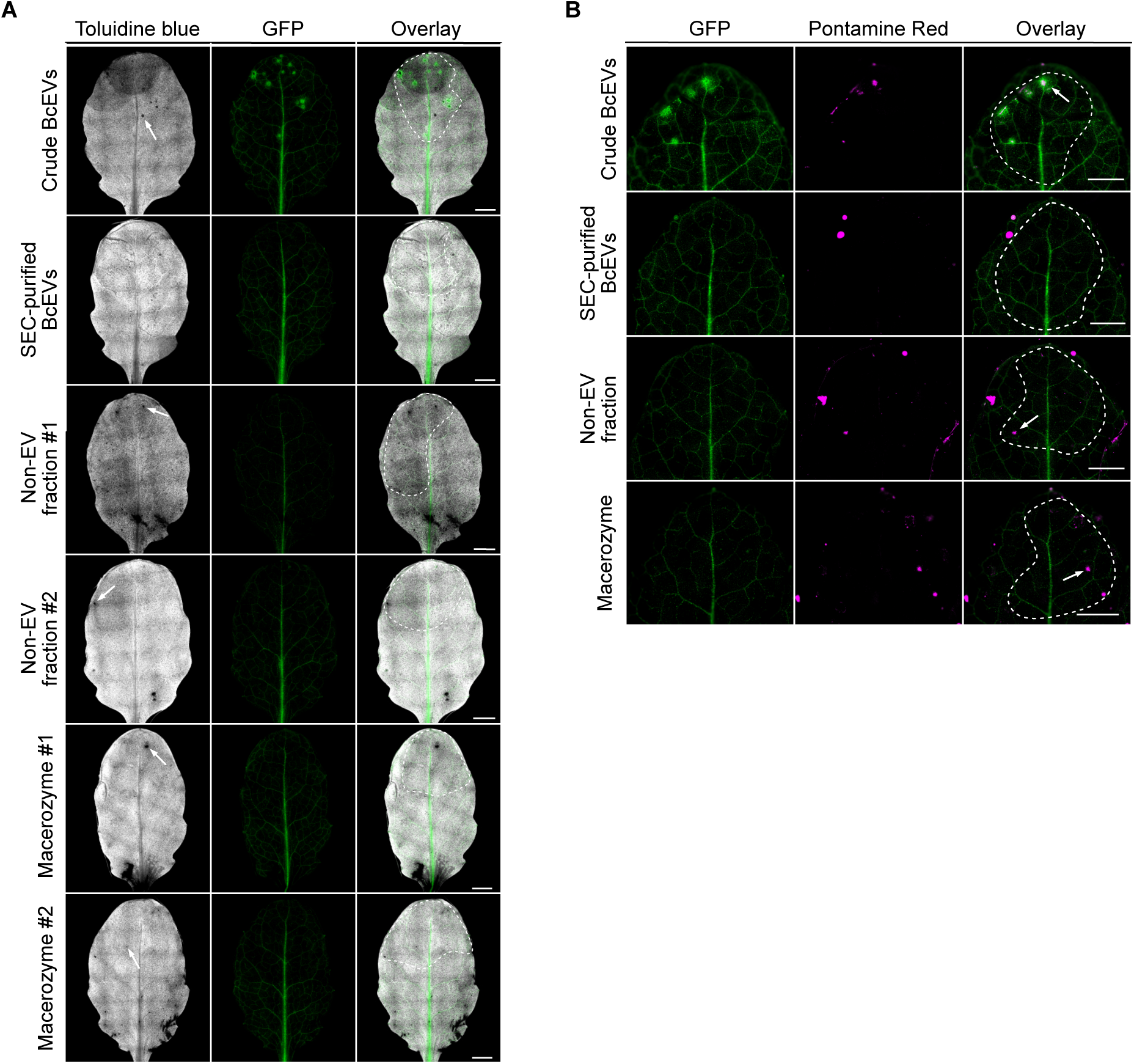
Replicates of cuticle permeability and cell wall staining assays after treatment with crude BcEVs, SEC-purified BcEVs, non-EV fraction or Macerozyme. A) GFP reporter plant leaves were drop-inoculated either with crude BcEVs, SEC-purified BcEVs, non-EV fraction and Macerozyme. Leaves were stained with toluidine blue and imaged in blue spectrum excitation and GFP channels. White arrows indicated toluidine staining signals. Scale bars represent 2 mm. B) GFP reporter plant leaves were drop-inoculated either crude BcEV, SEC-purified BcEV samples, non-EV fraction or Macerozyme. Leaves were stained with Pontamine fast scarlet 4B and imaged by fluorescence microscopy. Dotted circles indicate areas of treatment and white arrows indicated Pontamine signals. Scale bars represent 1 mm.

**Figure S18:**
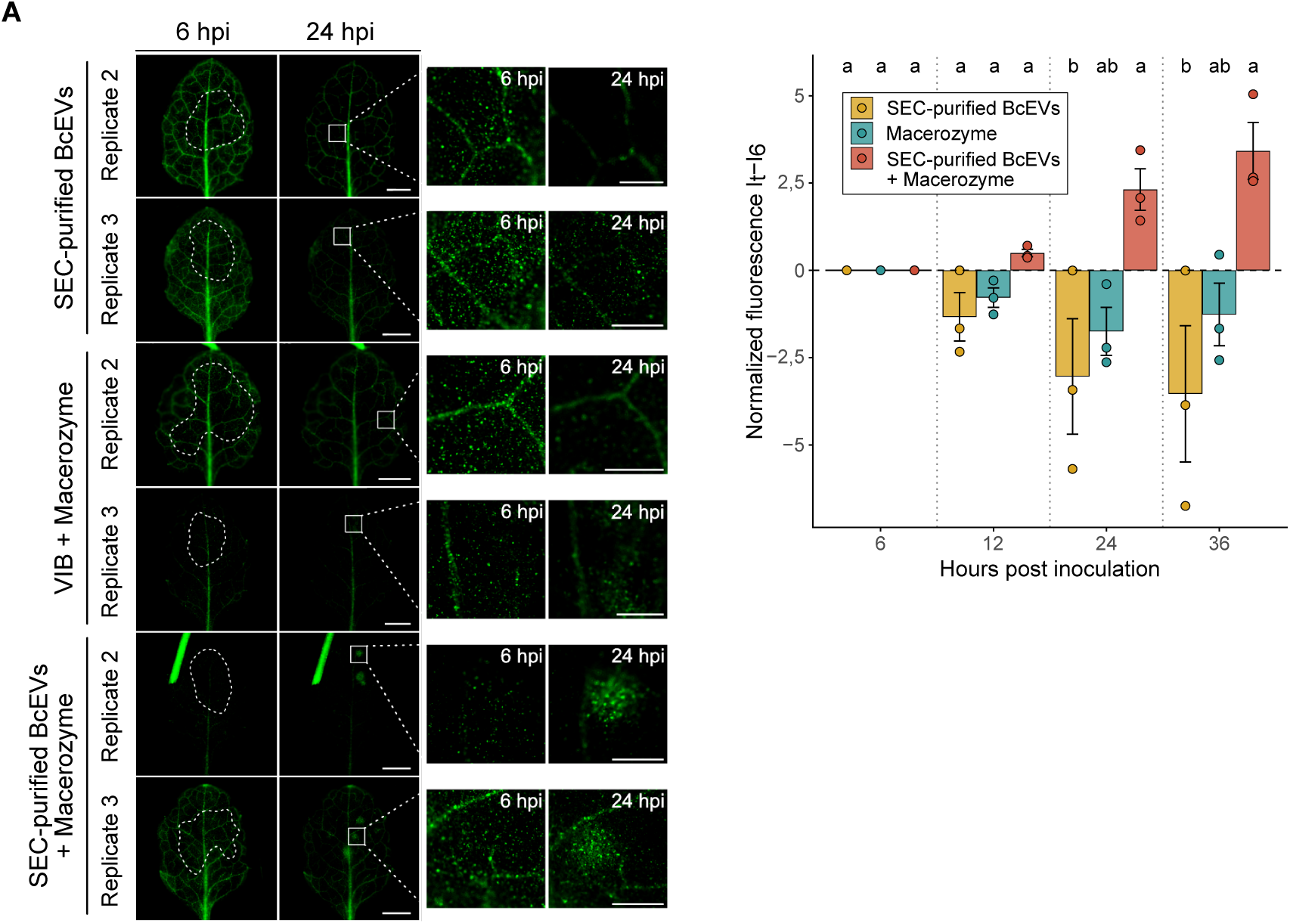
Replicates of Macerozyme treatment with SEC-purified BcEVs. A) Replicates of GFP reporter plants treated with SEC-purified BcEVs complemented with Macerozyme, according to Fig.4A. Dashed white lines outline the area of BcEV treatment; the white square at 24 hpi marks a representative site of GFP activation, shown magnified in the adjacent panels. Scale bars represent 2 mm. Normalized GFP fluorescence intensity (It – I6) at selected time points (6, 12, 24, and 36 hpi), corresponding to the data shown in Fig.4A and Fig.S18A. Bar graphs represent mean ± SEM of three biological replicates. Statistical significance was determined by pairwise ANOVA followed by Tukey’s HSD post hoc test at each time point; treatments not sharing the same letter are significantly different (p < 0.05).

**Figure S19:**
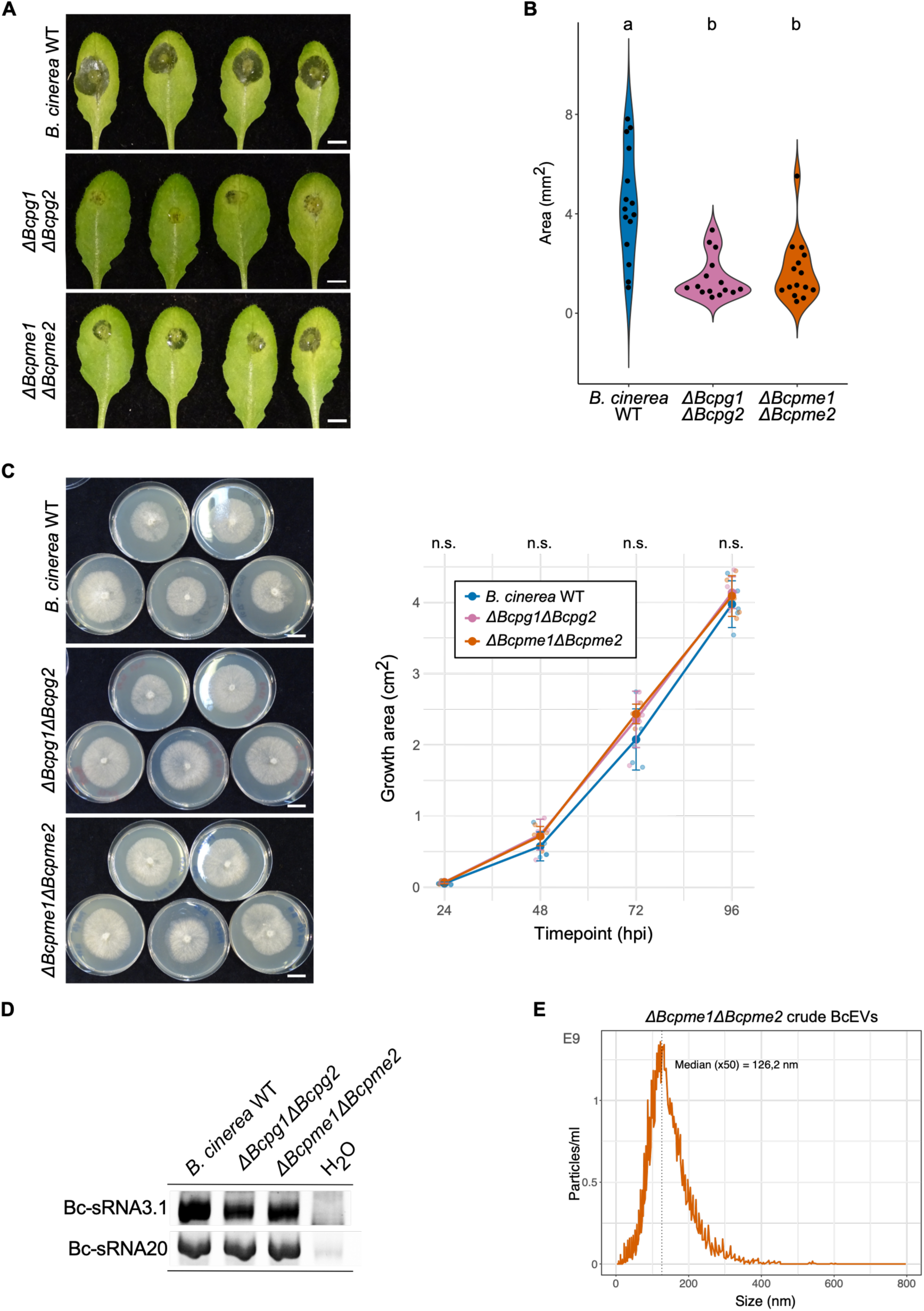
Infection assay and crude BcEV analysis of *B. cinerea ΔBcpme1ΔBcpme2* and *ΔBcpg1ΔBcpg2* mutants. A) Infection assay of *A. thaliana* leaves with *B. cinerea* WT, *ΔBcpg1ΔBcpg2* and *ΔBcpme1ΔBcpme2* mutants. Images shown infection at 72 hpi. Scale bars represent 5 mm. B) Measurement of infection lesion area induced by *B. cinerea* WT, *ΔBcpg1ΔBcpg2* and *ΔBcpme1ΔBcpme2* mutants. Letters indicate significant differences in lesion size using pairwise ANOVA with Tukey HSD p<0.05. Infection assays were repeated in 2 independent experiments with comparable results. C) Agar plate growth assay with *B. cinerea* WT, *ΔBcpg1ΔBcpg2* and *ΔBcpme1ΔBcpme2* mutant strains. Colony size on agar plates was measured in five independent cultures per strain every 24 hours. No statistical difference was determined when using pairwise ANOVA followed by Tukey’s HSD post hoc test at each time point. D) Stem-loop RT-PCR of Bc-sRNA3.1 and Bc-sRNA20 in *B. cinerea* WT *ΔBcpg1ΔBcpg2* and *ΔBcpme1ΔBcpme2* mutants. E) NTA plots of crude BcEVs isolated from *B. cinerea* WT and *ΔBcpme1ΔBcpme2* mutant. NTA measurements were repeated in 2 independent experiments with comparable results.

**Figure S20:**
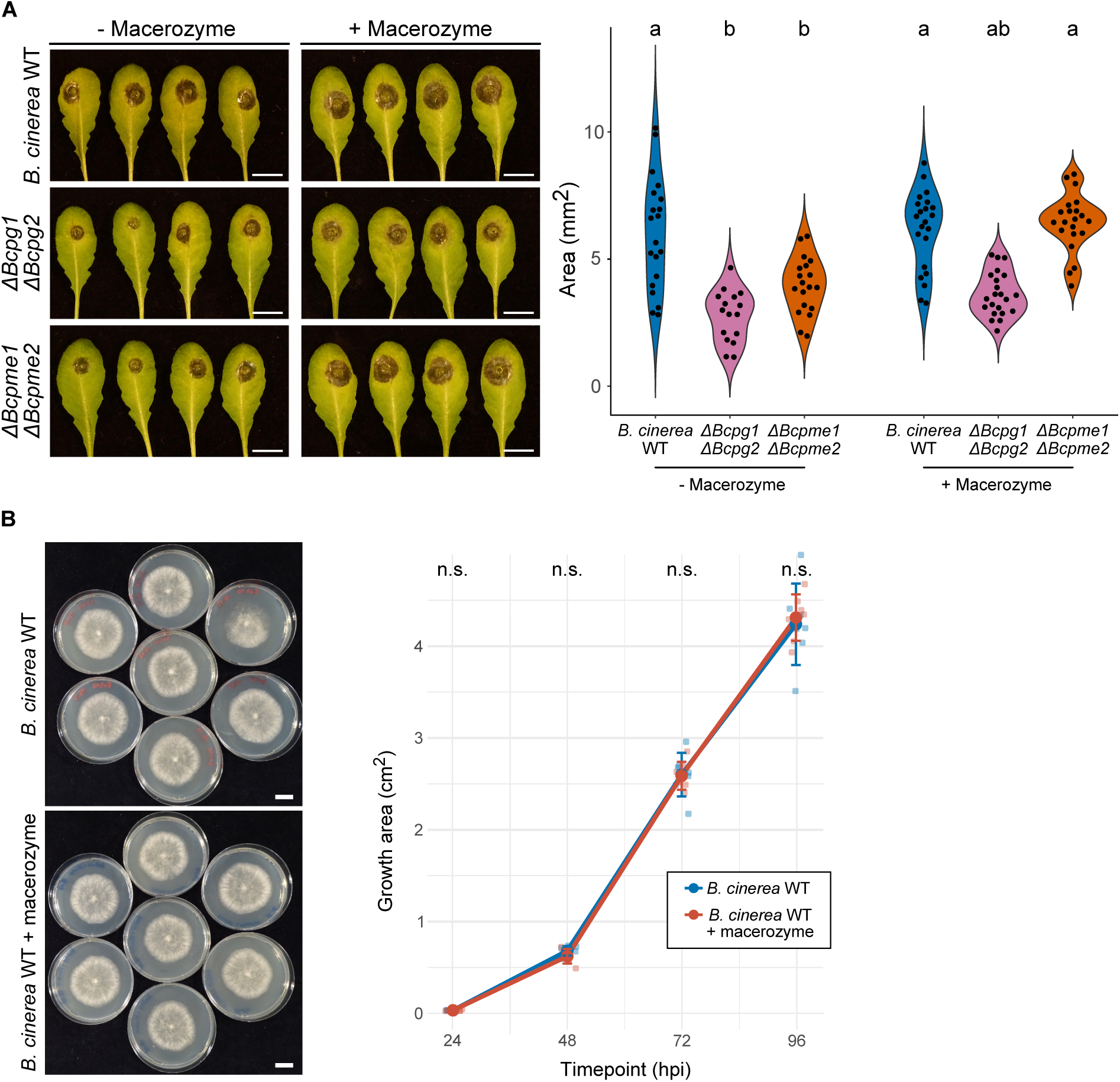
Macerozyme rescues infection of the *ΔBcpme1ΔBcpme2* mutant strain. A) Infection assay of *A. thaliana* leaves with *B. cinerea* WT, *ΔBcpg1ΔBcpg2* and *ΔBcpme1ΔBcpme2* mutants with the addition of Macerozyme. Images shown infection at 72 hpi. Scale bars represent 1 cm. Statistical significance was determined by pairwise ANOVA followed by Tukey’s HSD post hoc test at each time point; treatments not sharing the same letter are significantly different (p < 0.05). B) Agar plate growth assay with *B. cinerea* WT with the addition of Macerozyme. Colony size on agar plates was measured in seven independent cultures per strain every 24 hours. Statistical significance was determined by pairwise ANOVA followed by Tukey’s HSD post hoc test at each time point.

**Figure S21:**
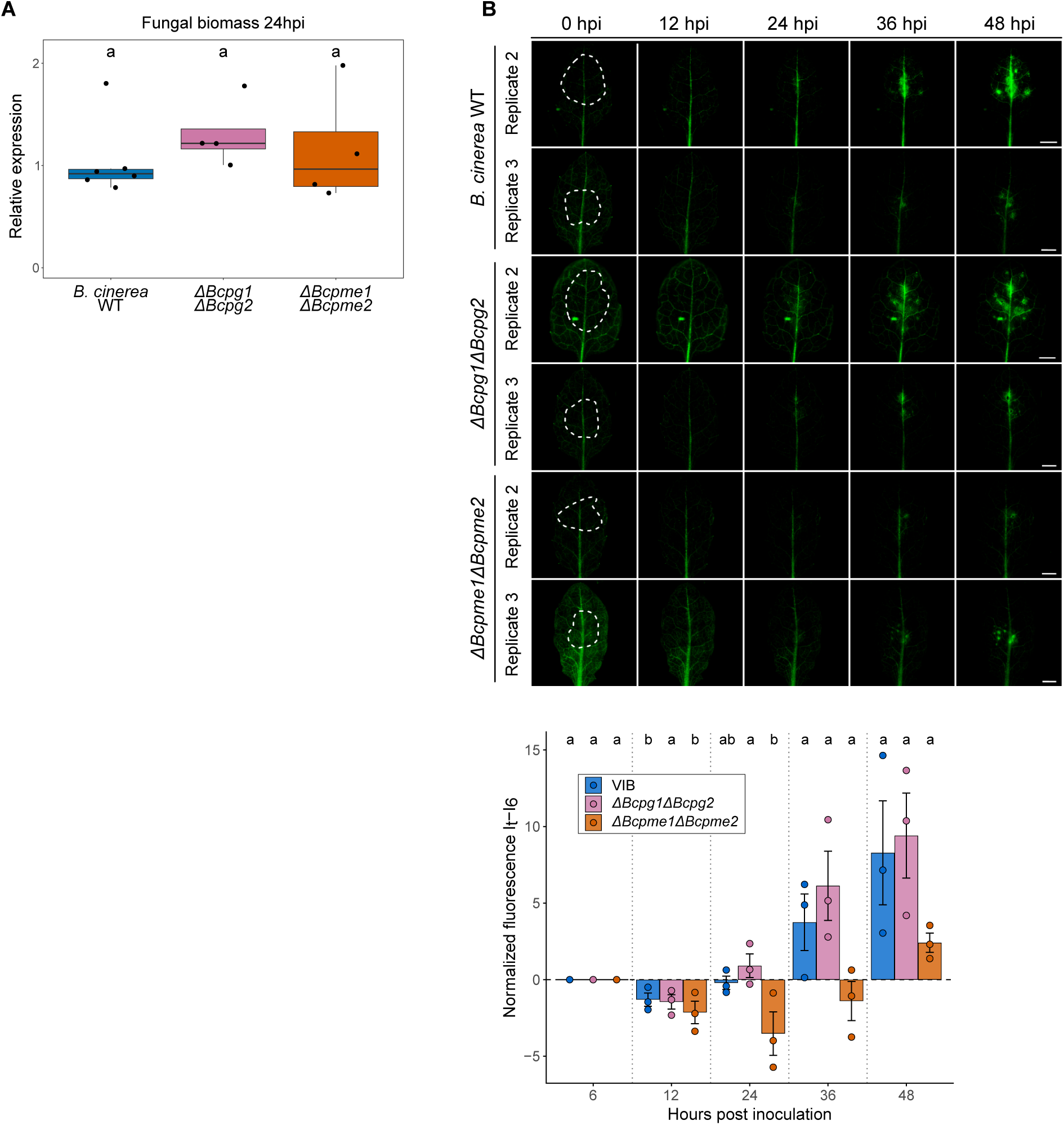
Replication of GFP reporter plants inoculated with *B. cinerea ΔBcpg1ΔBcpg2* and *ΔBcpme1ΔBcpme2* mutants. A) *B. cinerea* genomic DNA content relative the *A. thaliana* DNA in infected GFP reporter plants at 24 hpi measured by qPCR. *B. cinerea BcTubulin* and *A. thaliana AtActin* primers were used to quantify genomic DNA. Each data point represents a biological replicate. Letters indicate no significant difference according to ANOVA test with p<0.05. B) Replicate of GFP reporter plants infected with *B. cinerea* WT or *ΔBcpme1ΔBcpme2*, according to Fig.4B. Dotted lines indicate areas of treatment. Scale bars represent 2 mm. Normalized GFP fluorescence intensity (It – I6) at selected time points (6, 12, 24, and 36 hpi), corresponding to the data shown in Fig.4B and Fig.S21B. Bar graphs represent mean ± SEM of three biological replicates. Statistical significance was determined by pairwise ANOVA followed by Tukey’s HSD post hoc test at each time point; treatments not sharing the same letter are significantly different (p < 0.05).

**Figure S22:**
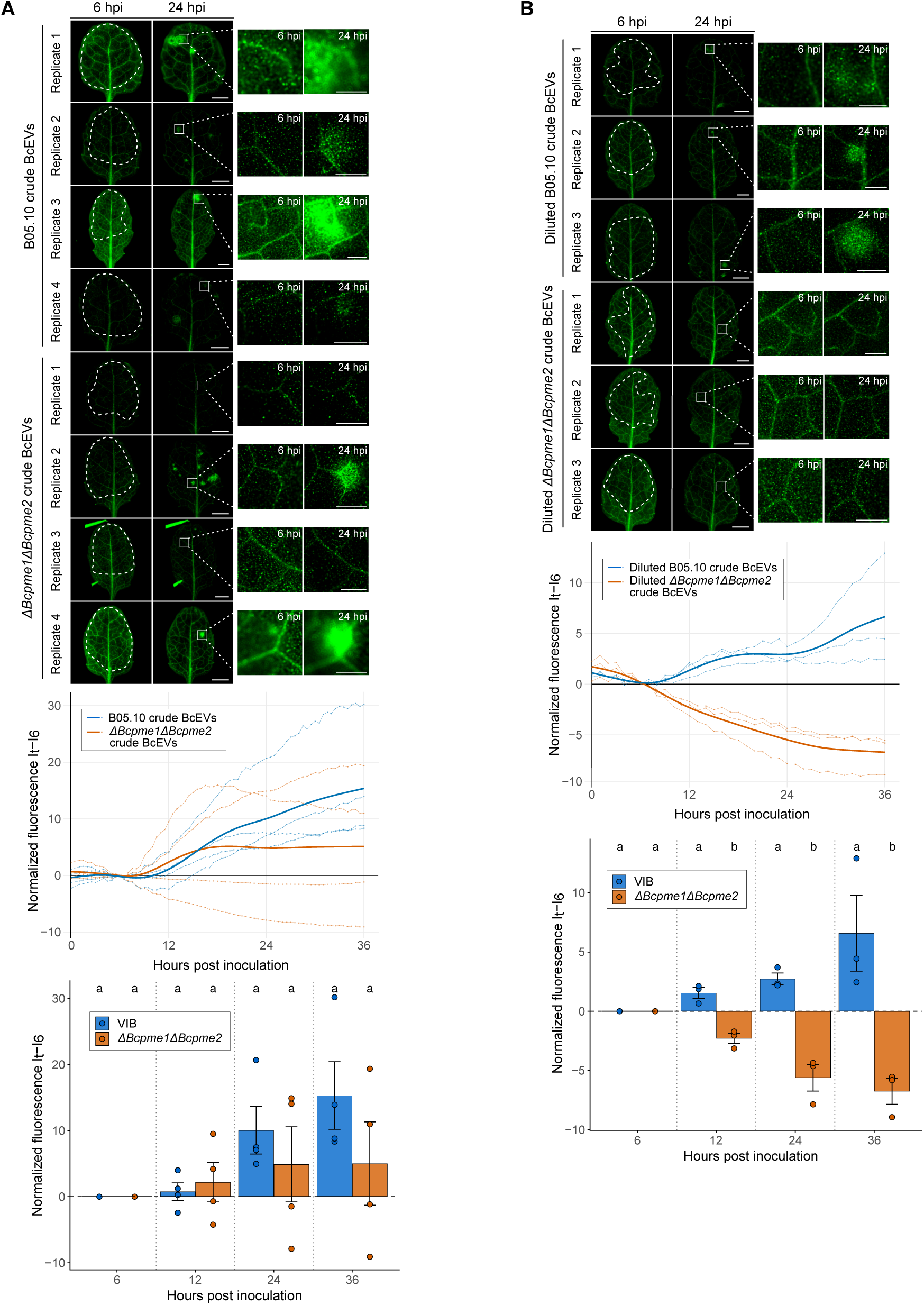
GFP reporter plants treated with crude BcEVs from *B. cinerea ΔBcpme1ΔBcpme2*. A) GFP reporter plants were treated with 5 µl crude BcEVs (2,4×10^14^ particles/ml) of WT or of the *ΔBcpme1ΔBcpme2* mutant. Time-course fluorescence microscopy was carried out from 0 - 36 hpi. Dashed white lines outline the area of BcEV treatment; the white square at 24 hpi marks a representative site of GFP activation, shown magnified in the adjacent panels. 2 out of 4 leaves treated with *ΔBcpme1ΔBcpme2* BcEVs showed GFP activation. Scale bars represent 2 mm. Normalized GFP fluorescence intensity (It – I6) at selected time points (6, 12, 24, and 36 hpi), corresponding to the data shown in Fig.S22A. Bars represent mean ± SEM of three biological replicates. Statistical significance was determined by pairwise ANOVA followed by Tukey’s HSD post hoc test at each time point. B) GFP reporter plants were treated with 5µl 1:3 diluted crude BcEVs of WT or of the *ΔBcpme1ΔBcpme2* mutant. Time-course fluorescence microscopy was carried out from 0 - 36 hpi. Dashed white lines outline the area of BcEV treatment; the white square at 24 hpi marks a representative site of GFP activation, shown magnified in the adjacent panels. Scale bars represent 2 mm. Normalized GFP fluorescence intensity (It – I6) at selected time points (6, 12, 24, and 36 hpi), corresponding to the data shown in Fig.S22B. Bar graphs represent mean ± SEM of three biological replicates. Statistical significance was determined by pairwise ANOVA followed by Tukey’s HSD post hoc test at each time point; treatments not sharing the same letter are significantly different (p < 0.05).

**Figure S23:**
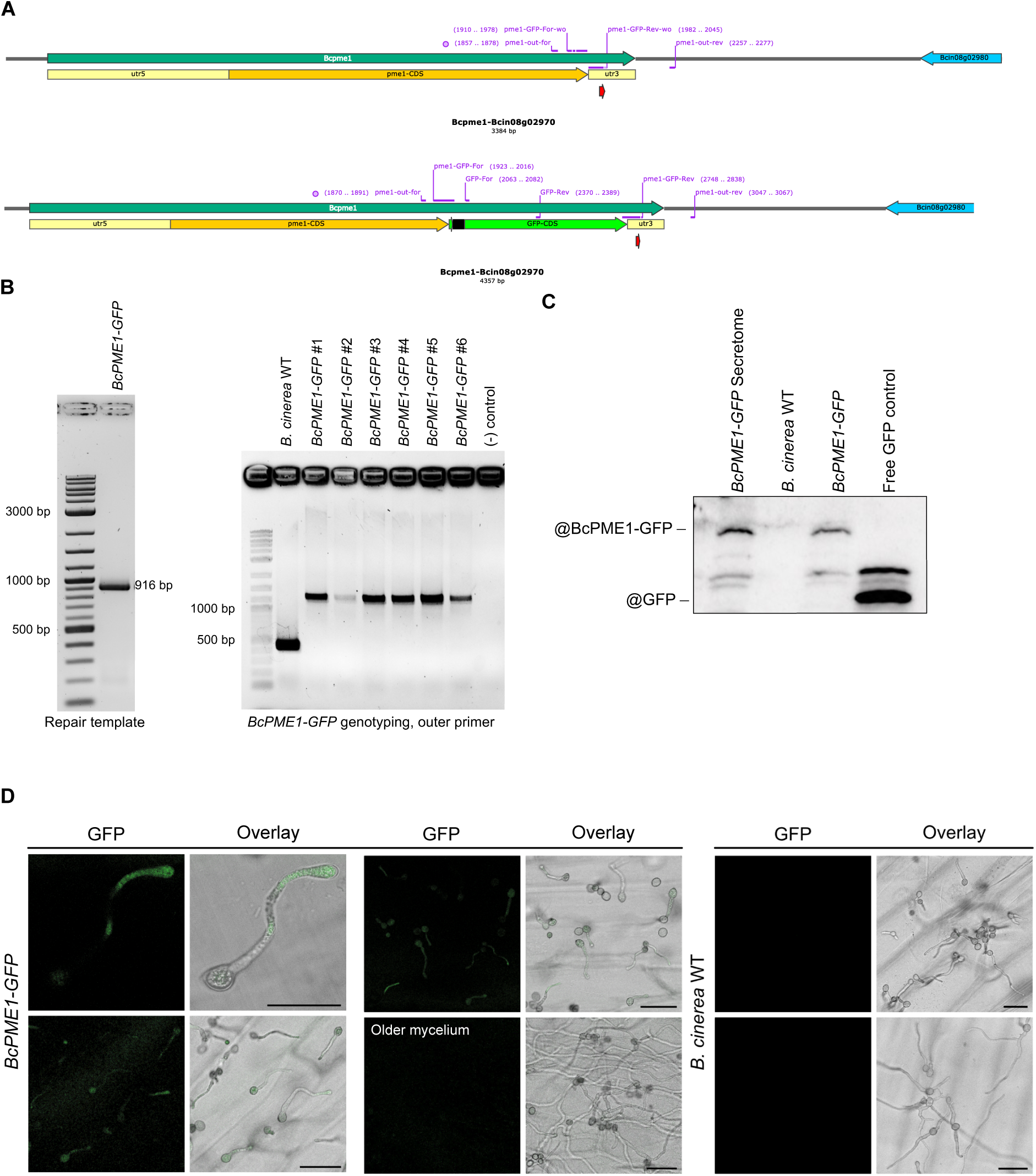
BcPME1-GFP is expressed in infecting *B. cinerea* hyphae. A) Schematic view of the Cas9-mediated knock-in of the GFP transgene into the C-terminal position of the native *Bcpme1* locus. Primer binding sites used for genotyping and sequencing are indicated in purple. Red arrowhead marks the guide RNA target site used for Cas9-mediated insertion. B) PCR validation of the *BcPME1-GFP* fusion in isolated fungal transformants using genomic DNA. C) Immunoblot analysis of BcPME1-GFP in fungal total protein extracts of mycelium and culture supernatant (secretome) using an anti-GFP antibody. Free GFP and *B. cinerea* WT extracts were used as controls. D) Confocal microscopy of *B. cinerea* WT and a BcPME1-GFP expressing strain during infection. GFP and corresponding brightfield overlay images are shown for young hyphae, older mycelium, and *B. cinerea* WT control. Scale bars represent 25 µm (top left) and 50 µm for all other images.

**Figure S24:**
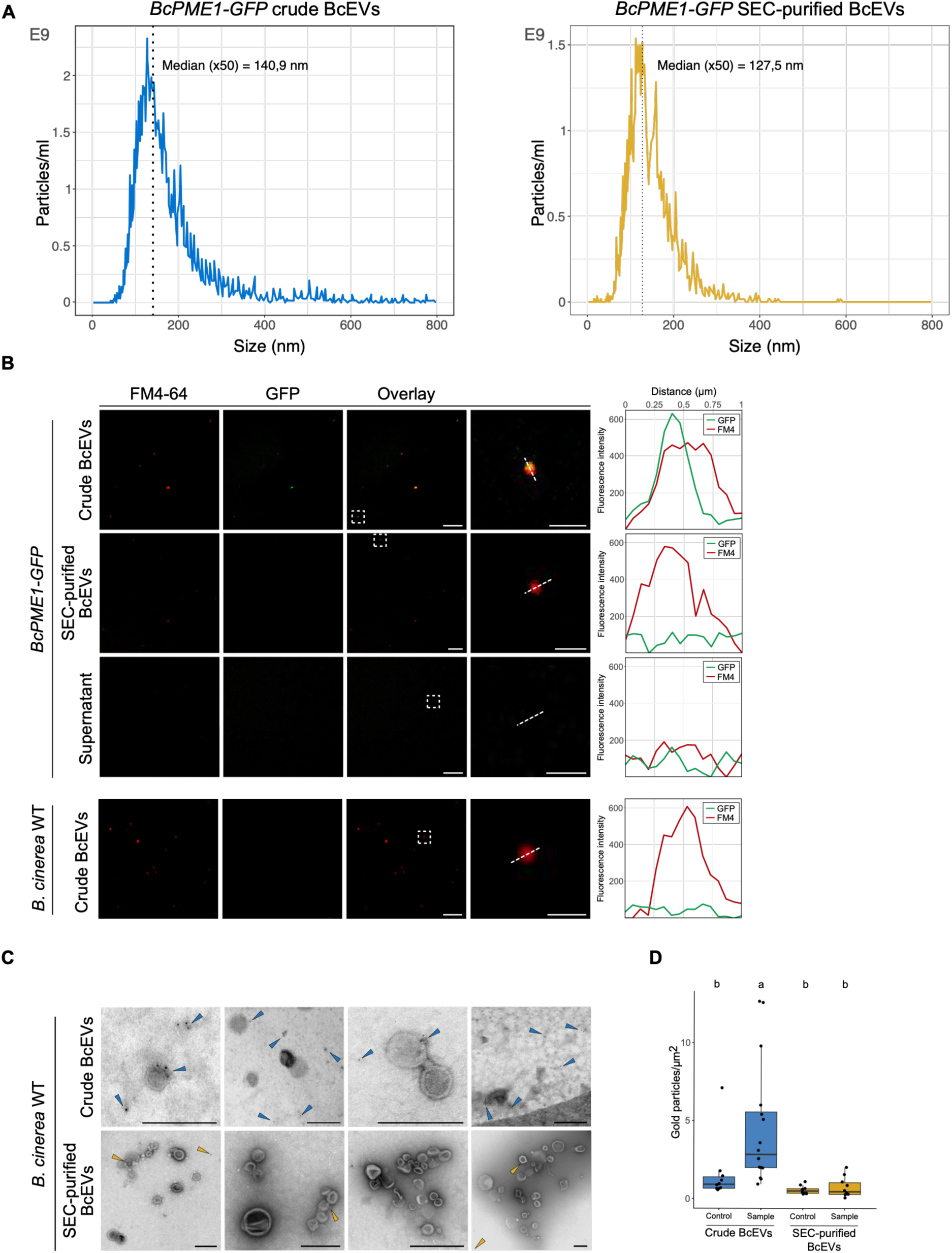
BcPME1-GFP associates with crude BcEVs. A) NTA blot of crude BcEVs and SEC-purified BcEVs collected from the *B. cinerea BcPME1-GFP* strain. B) Crude BcEVs, SEC-purified BcEVs, and the supernatant fraction from the *BcPME1-GFP* strain, as well as crude BcEVs from *B. cinerea* WT, were stained with the lipophilic dye FM4-64. Overlay panels show colocalization of GFP and FM4-64 signal; regions inside the dashed squares are magnified in the adjacent panel. Fluorescence intensity profiles were recorded along the indicated dashed line, showing spatial overlap of GFP and FM4-64 signal. Scale bars represent 1 µm for the magnified images and 5 µm for all the other images. C) Immunogold labeling and TEM analysis of *B. cinerea* WT crude BcEVs and SEC-purified BcEVs using an anti-Pectinase antibody. Arrowheads indicate gold particles. Scale bars represent 250 nm. D) Gold particles per µm² were counted in 11 TEM images of both crude BcEVs and SEC-purified BcEVs. Control samples were incubated only with secondary antibody. Statistical significance was determined by pairwise ANOVA followed by Tukey’s HSD post hoc test; groups not sharing the same letter are significantly different (p < 0.05).

**Figure S25:**
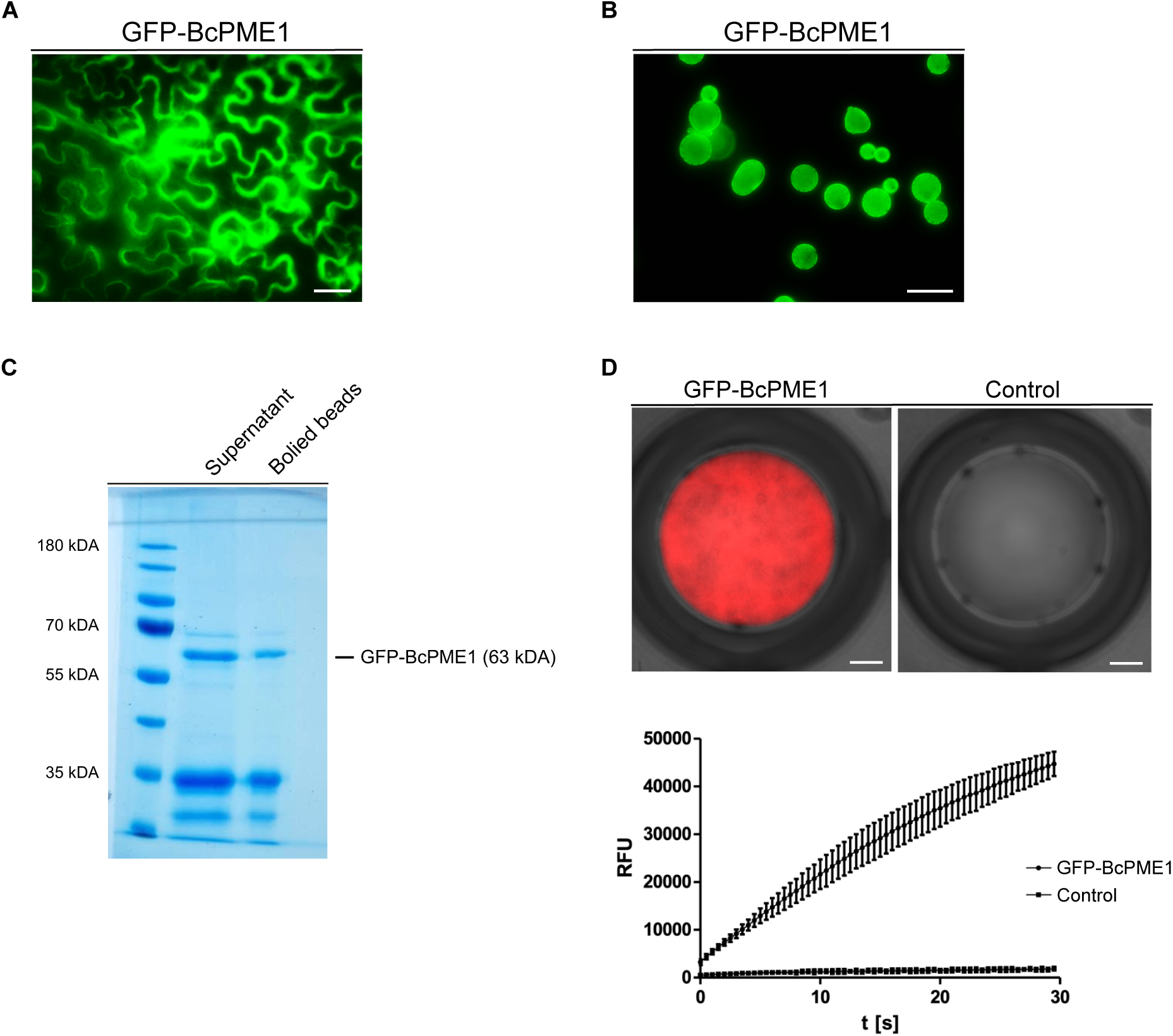
Heterologous GFP-BcPME1 expression, purification and enzymatic activity. A) Transient GFP-BcPME1 expression in tobacco leaves was achieved by *Agrobacterium*-mediated transformation. Scale bar represents 40 µm. B) GFP-BcPME1 purification using anti-GFP magnetic beads. Scale bar represents 50 µm. C) Purified BcPME1 was separated on an SDS-PAGE and stained by Coomassie Brilliant Blue. Anti-GFP magnetic beads were boiled in 2x SDS buffer and supernatant and boiled beads were loaded onto the SDS-PAGE. D) BcPME1 activity assay using a modified Ampliflu-Red assay. Scale bars represent 1 mm.

**Figure S26:**
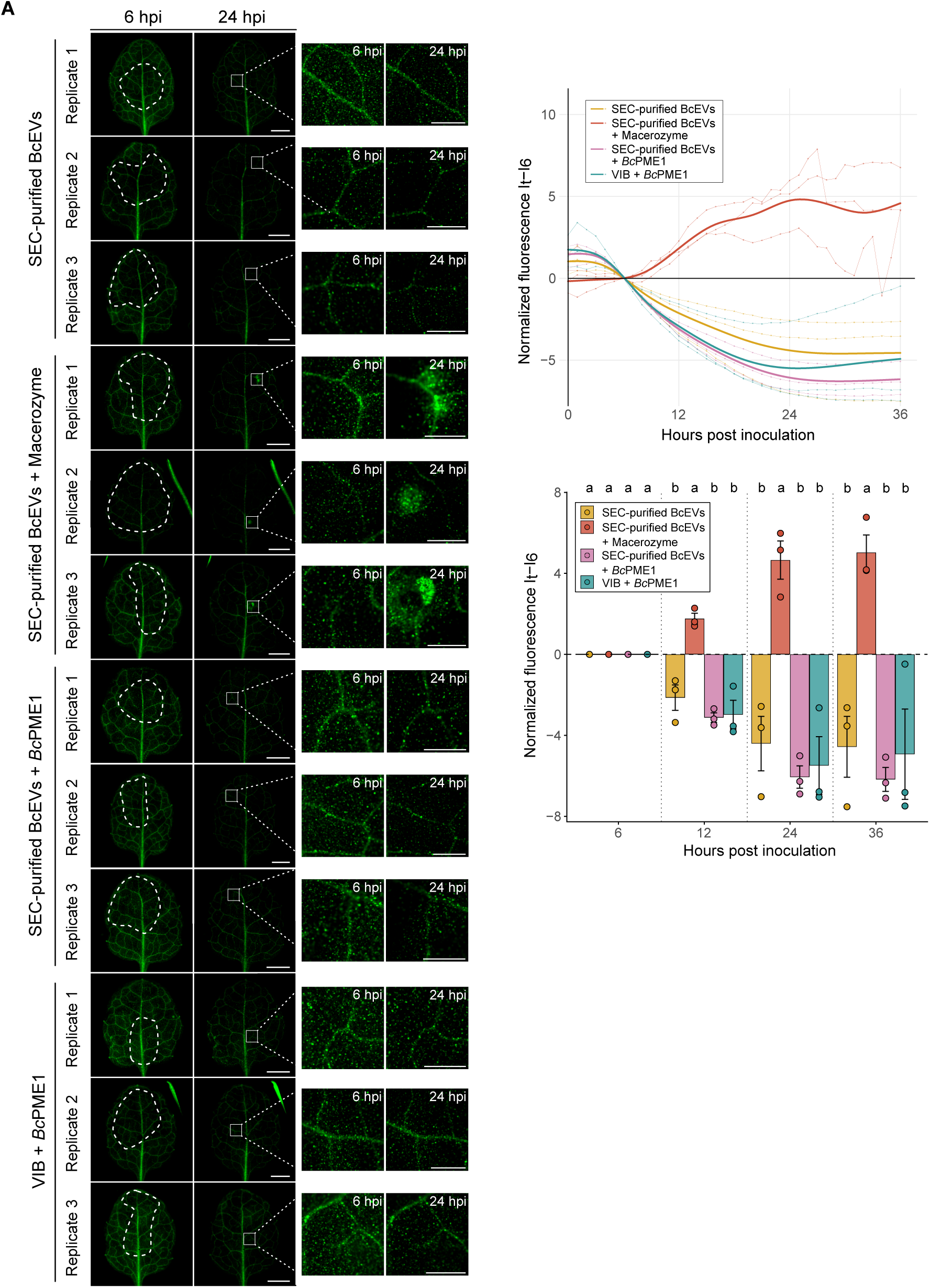
Complementation assay of SEC-purified BcEVs with BcPME1. A) GFP reporter plants were treated with 5µl SEC-purified BcEVs (4,9×10^13^ particles/ml) mixed with purified GFP-BcPME1 in a 1:1 ratio. SEC-purified BcEVs complemented with Macerozyme were used as positive control. Dashed white lines outline the area of BcEV treatment; the white square at 24 hpi marks a representative site of GFP activation, shown magnified in the adjacent panels. Scale bars represent 2 mm. Normalized GFP fluorescence intensity (It – I6) at selected time points (6, 12, 24, and 36 hpi), corresponding to the data shown in Fig.S26A. Bar graphs represent mean ± SEM of three biological replicates. Statistical significance was determined by pairwise ANOVA followed by Tukey’s HSD post hoc test at each time point; treatments not sharing the same letter are significantly different (p < 0.05).

**Figure S27:**
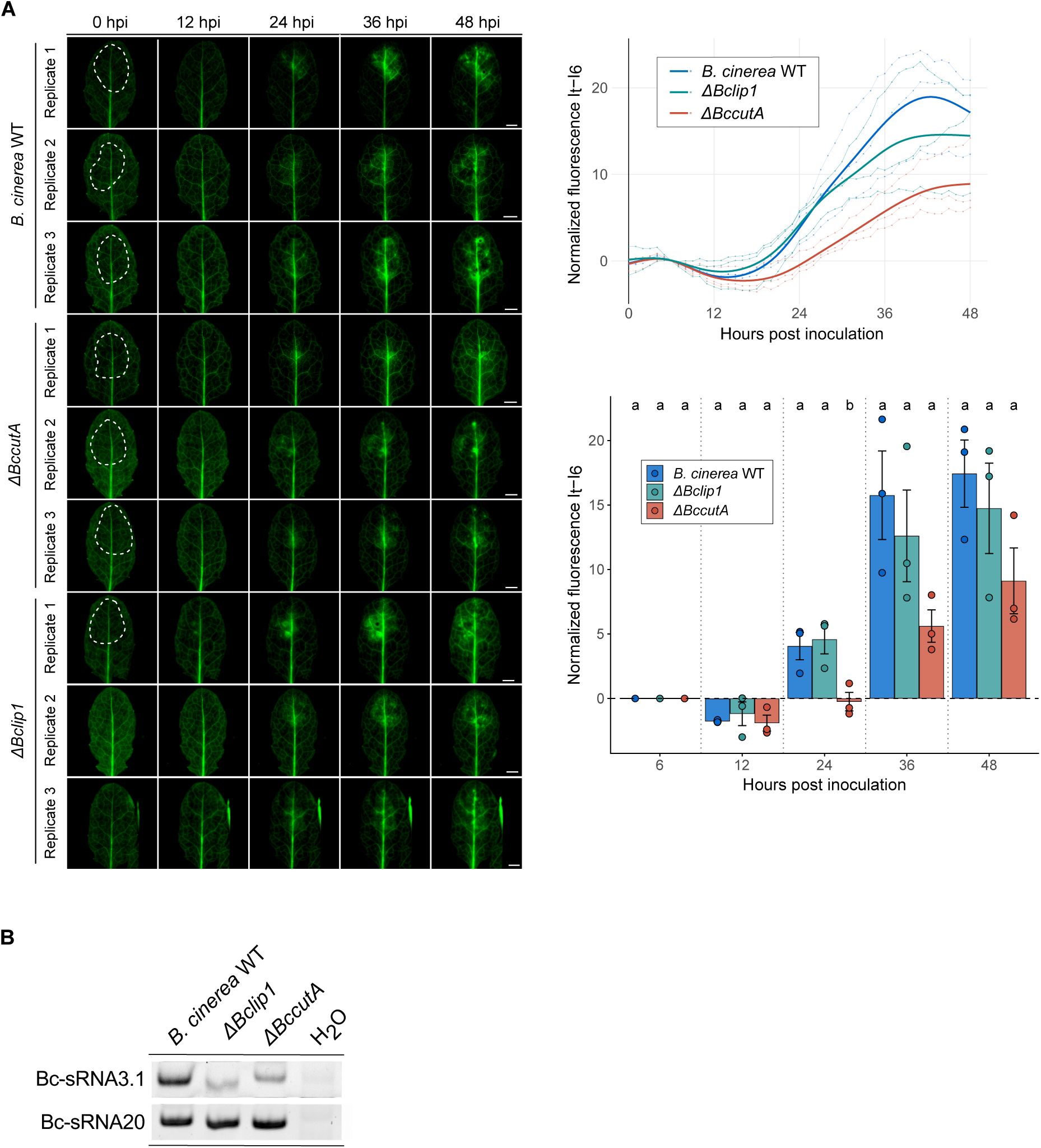
GFP reporter plant assay with *B. cinerea ΔBclip1* and *ΔBccutA* mutants. A) GFP reporter plants infected with *B. cinerea* WT, *ΔBclip1* and *ΔBccutA*. Dotted lines indicate areas of treatment. Scale bars represent 2 mm. Normalized GFP fluorescence intensity (It – I6h) at selected time points (6, 12, 24, and 36 hpi), corresponding to the data shown in Fig.S27A. Bar graphs represent mean ± SEM of three biological replicates. Statistical significance was determined by pairwise ANOVA followed by Tukey’s HSD post hoc test at each time point; treatments not sharing the same letter are significantly different (p < 0.05). B) Stem-loop RT-PCR of Bc-sRNA3.1 and Bc-sRNA20 in *B. cinerea* WT, *ΔBclip1* and *ΔBccutA* mutants.

**Figure S28:**
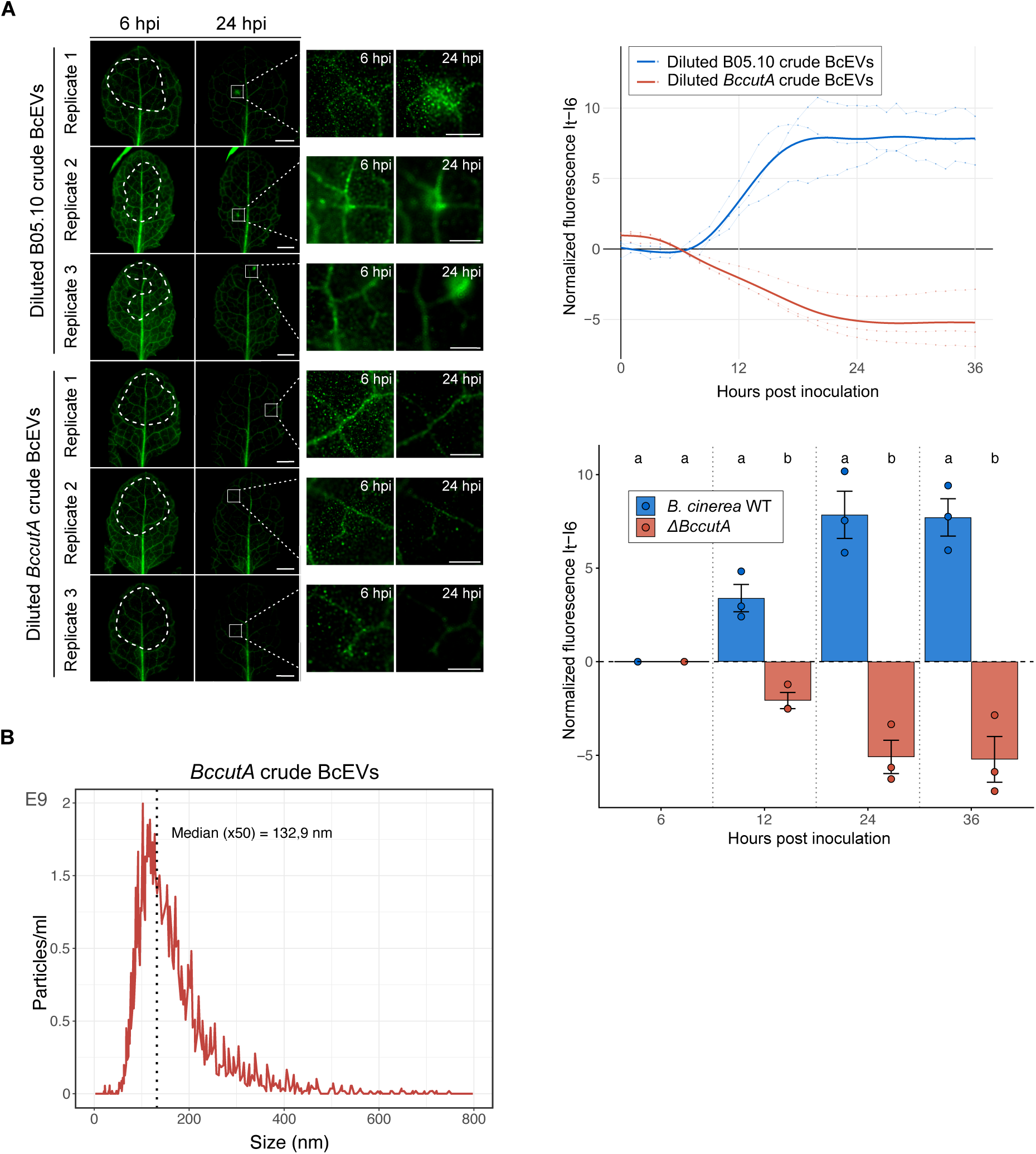
GFP reporter plants treated with crude BcEVs from *B. cinerea ΔBccutA* mutant. A) GFP reporter plants were treated with 5µl 1:3 diluted crude BcEVs (1,3×10^14^ particles/ml) of WT or of the *ΔBccutA* mutant. Time-course fluorescence microscopy was carried out from 0 - 36 hpi. Dashed white lines outline the area of BcEV treatment; the white square at 24 hpi marks a representative site of GFP activation, shown magnified in the adjacent panels. Scale bars represent 2 mm. Normalized GFP fluorescence intensity (It – I6) at selected time points (6, 12, 24, and 36 hpi), corresponding to the data shown in Fig.S28A. Bar graphs represent mean ± SEM of three biological replicates. Statistical significance was determined by pairwise ANOVA followed by Tukey’s HSD post hoc test at each time point; treatments not sharing the same letter are significantly different (p < 0.05). B) NTA blot of crude BcEVs of the *ΔBccutA* mutant.

**Figure S29:**
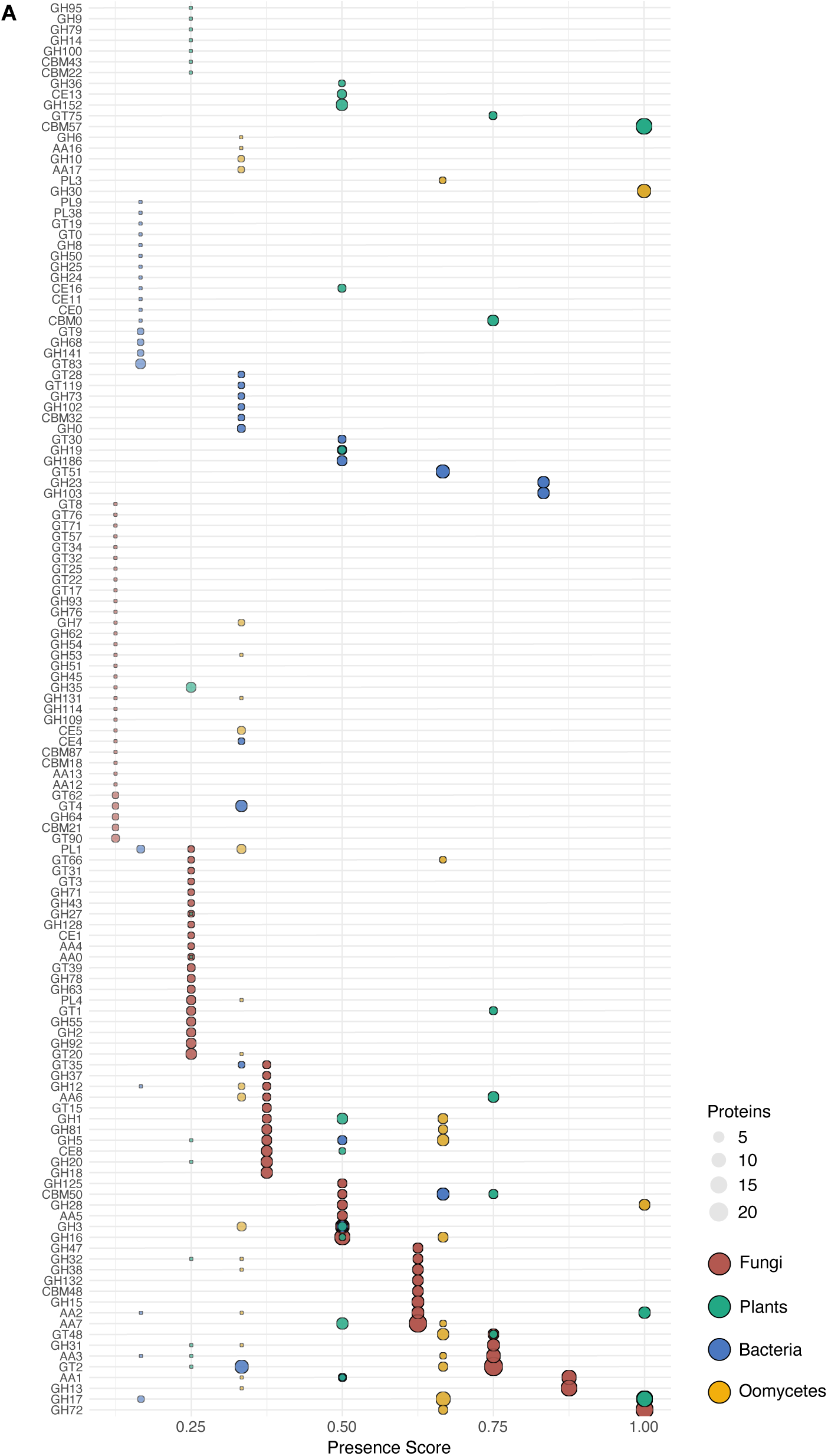
Presence matrix of all CAZymes identified in BcEV samples among different kingdoms. A) Analysis was run on publicly available EV protein datasets (Tab.S4, Tab S5).

